# Early Detection of Sudden Transitions in Notch signaling

**DOI:** 10.64898/2026.02.01.703083

**Authors:** Baishakhi Tikader, Sukanta Sarkar, Sudipta Kumar Sinha, Herbert Levine, Mohit Kumar Jolly, Partha Sharathi Dutta

**Affiliations:** Department of Bioengineering Indian Institute of Science, Bengaluru, India 560012; Department of Mathematics Indian Institute of Technology Ropar, Rupnagar, Punjab 140 001 India; Department of Chemistry Indian Institute of Technology Ropar, Rupnagar, Punjab 140 001 India; Department of Physics and Department of Bioengineering Northeastern University, Boston, MA 02115, USA; Center for Theoretical Biological Physics Northeastern University, Boston, MA 02115, USA

**Keywords:** Notch-Delta-Jagged pathway, critical transitions, flickering, early warning signals, rate of forcing, demographic noise

## Abstract

Identifying sudden transitions during phenotypic decision-making of complex biological systems can be crucial for our ability to control a cellular state. Yet, prior determination of these sudden transitions or tipping points remains challenging, as biological systems often exhibit only subtle early changes, which are often masked by inherent noise or rapid transition dynamics. Using Notch signaling as a model, we systematically analyze dynamical transitions in Notch-Delta (ND), Notch-Delta-Jagged (NDJ), and Fringe-mediated NDJ systems for both one and two-cell contexts. In the one-cell ND system, critical slowing down (CSD)-based early warning signals (EWSs) reliably capture transitions between sender (*S*) and receiver (*R*) states and remain robust to variation in forcing rate. We further find that flickering is a precursor to transitions in one-cell NDJ system. In contrast, flickering does not occur in the two-cell Notch model due to the presence of a supercritical bifurcation. Our analysis also offers insight into how NICD (Notch Intracellular Domain)-driven and Fringe-mediated asymmetries, along with the strength of external signals, control the emergence of flickering. Overall, this study identifies sudden transitions in Notch signaling under demographic noise and can be extended to other noisy biological systems, with potential applications in drug development and targeted therapeutic interventions.

## 2 Introduction

The ability of a biological system to switch between multiple stable states often has a crucial role in controlling various life processes, such as cell differentiation (Duddu et al 2020, Ghaffarizadeh et al 2014), cellular plasticity (Jia et al 2019), and cellular reprogramming (Bargaje et al 2017, MacArthur et al 2009). The Notch signaling pathway is one of such fundamental intercellular communication systems, where the cell-cell coordination plays a crucial role during development, tissue homeostasis, and regeneration (Shi et al 2024, Zhou et al 2022, Siebel and Lendahl 2017, Gozlan and Sprinzak 2023, Artavanis-Tsakonas et al 1995, Mumm and Kopan 2000). This pathway functions through direct physical interactions between membrane-bound receptors (Notch) and ligands of the Delta-like and Jagged families expressed in neighboring cells (Jolly et al 2015, Bocci et al 2019, Ehebauer et al 2006). Notch-Delta interactions typically drive lateral inhibition, causing neighboring cells to adopt opposite fates, as is observed during Drosophila neurogenesis and vertebrate inner ear cell patterning (Campos-Ortega et al 1993, Kelley 2006). In contrast, the Notch-Delta-Jagged system can generate lateral induction, where neighboring cells converge toward similar fates due to Jagged-mediated cooperative activation, as observed in processes such as endothelial tip/stalk cell coordination, epithelial-mesenchymal plasticity, and vascular smooth muscle differentiation (Suchting et al 2007, Lowell et al 2006). The combinatorial complexity arising from multiple receptor-ligand (R/L) variants, their post-translational modifications, and asymmetric ligand binding affinity to Notch allows Notch to generate diverse, context-specific fate decisions (D’souza et al 2008, Bray 2006, Nandagopal et al 2018). Furthermore, R/L binding strength is fine-tuned by Fringe-mediated glycosylation of Notch, which preferentially enhances Delta binding and suppresses Jagged binding affinity, thereby further shaping the patterning and stability of cell fate outcomes (Moloney et al 2000, Brückner et al 2000, Bocci et al 2020, Mukherjee and Levine 2023, Bray 2016).

Given the pivotal role of Notch in controlling cell-fate decisions, it is crucial to identify the sudden transitions in Notch-mediated cell-cell interaction dynamics. Transitions have been studied extensively in many natural systems, including ecology, and systems biology (Scheffer et al 2012, Liu et al 2014, Trefois et al 2015, van de Leemput et al 2014, Kianercy et al 2014). Sudden transitions from one asymptotically stable state to another upon crossing a critical threshold showing the critical slowing down (CSD) phenomenon as the system approaches a critical threshold or a tipping point (Scheffer et al 2009, Axelrod et al 2015, Sharma et al 2016, Li et al 2018) - is a characteristic feature of various bi-stable/multistable systems, as investigated in many naturally occurring or synthetically constructed gene regulatory networks. Although recent studies have established to find the signs of critical transitions (Axelrod et al 2015, Sharma et al 2016) in intracellular processes involved in cell division and in the onset of cancer metastasis (Yang et al 2018, Sarkar et al 2019, Kashyap et al 2025), such analyses are largely absent for intercellular signaling pathways. Specifically, identifying critical transitions in the Notch pathway (Boareto et al 2016, Bocci et al 2017) has not yet been investigated.

It is evident that there are intrinsic fluctuations in biological processes, impacting gene regulation to transcription to ligand production and receptor activation (Munsky et al 2009, Selvam 2017, Paulsson 2004, Bar-Even et al 2006, Munsky et al 2012, Tsimring 2014). Such stochasticity can significantly influence phenotypic outcomes (Pilpel 2011, Balázsi et al 2011). For example, intrinsic noise can modulate oscillatory behaviors in neighboring cell populations communicating via Notch-Delta signaling (Baron and Galla 2019). Similarly, Notch signaling may be coupled to other oscillatory intracellular circuits (Singh et al 2021). These observations underscore the significance of investigating noise-induced transitions in systems where signaling relies on cell-cell contact.

Here, we first focus on identifying sudden transitions and the associated CSD in one and two-cell systems containing Notch-Delta and Notch-Delta-Jagged interactions. Then, we identify flickering as a new class of early warning signal for critical transitions operating in the Notch-Delta-Jagged system, representing the system’s ability to fluctuate between two alternative basins of attraction. By performing numerical simulations, we have shown that the balance between the relative production rates of the two ligands, Delta versus Jagged, can control the noisy behavior. In addition, under certain parameter regimes observed in the Notch–Delta–Jagged system, the presence of flickering can be captured by an effective stochastic potential. Finally, we showed how the interplay between the Fringe-mediated asymmetric effect on R/L binding and the variable expression of both ligands level by either altering production rate or external ligand-mediated signals determines the sensitivity of the flickering.

## 3 Mathematical Model

To explore the transitions and early warning signals in an intercellular context, we consider a model of the Notch signaling pathway proposed by several groups (Sprinzak et al 2010, Boareto et al 2015). The Notch signaling circuit operates through two distinct modes of interaction: cis and trans. In trans-interactions (also known as trans-activation), the Notch receptor (*N*) on a given cell engages with Jagged or Delta ligands (*J*_ext_ or *D*_ext_) presented on neighboring cells. This intercellular binding event triggers proteolytic cleavage of the receptor, releasing the Notch intracellular domain (NICD), denoted by *I*, which subsequently translocates to the nucleus and indirectly activates Notch and Jagged and represses Delta (Fig. 1A) and propagates the downstream signaling response. On the other hand, in the cis-interaction, also known as cis-inhibition, the receptor (*N*) of one cell interacts with either of the ligands within the same cell (*D* or *J*), causing the degradation of both interacting proteins. (Fig. 1A) These interactions have been described by the following deterministic equations (Boareto et al 2015):

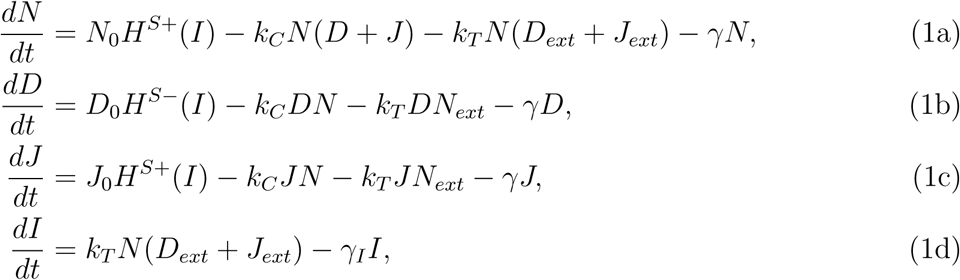

where *N, D, J*, and *I* represent Notch, Delta, Jagged, and NICD, respectively. *γ* represents the degradation rate of all three proteins (Notch, Delta, and Jagged), and *γ_I_* is the degradation rate of NICD. *k_C_* and *k_T_* are the strengths of cis-inhibition and trans-activation, respectively. *N_ext_*, *D_ext_*, and *J_ext_* represent the amount of ligands available for binding, which are present on the membrane surfaces of neighbouring cells. *N*_0_, *D*_0_, and *J*_0_ are the production rates of Notch, Delta, and Jagged, respectively. Here, the effect of NICD (*I*) on Notch, Delta, and Jagged has been incorporated by introducing a shifted Hill function (*H^S^*(*I, λ*)) for the production rates of these proteins. *H^S^*(*I, λ*) = *H^−^*(*I*) + *λH*^+^(*I*), 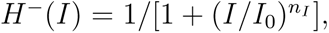 *H*^+^(*I*) = 1 *− H^−^*(*I*). *λ* represents the fold change in the production rate, and for activation *λ >* 1, whereas for inhibition *λ <* 1. The steepness of the Hill function is determined by the power *n_I_*. For more details of the model derivation (Lu et al 2013) see **SI Text S1.1.1**.

**Figure 1.**
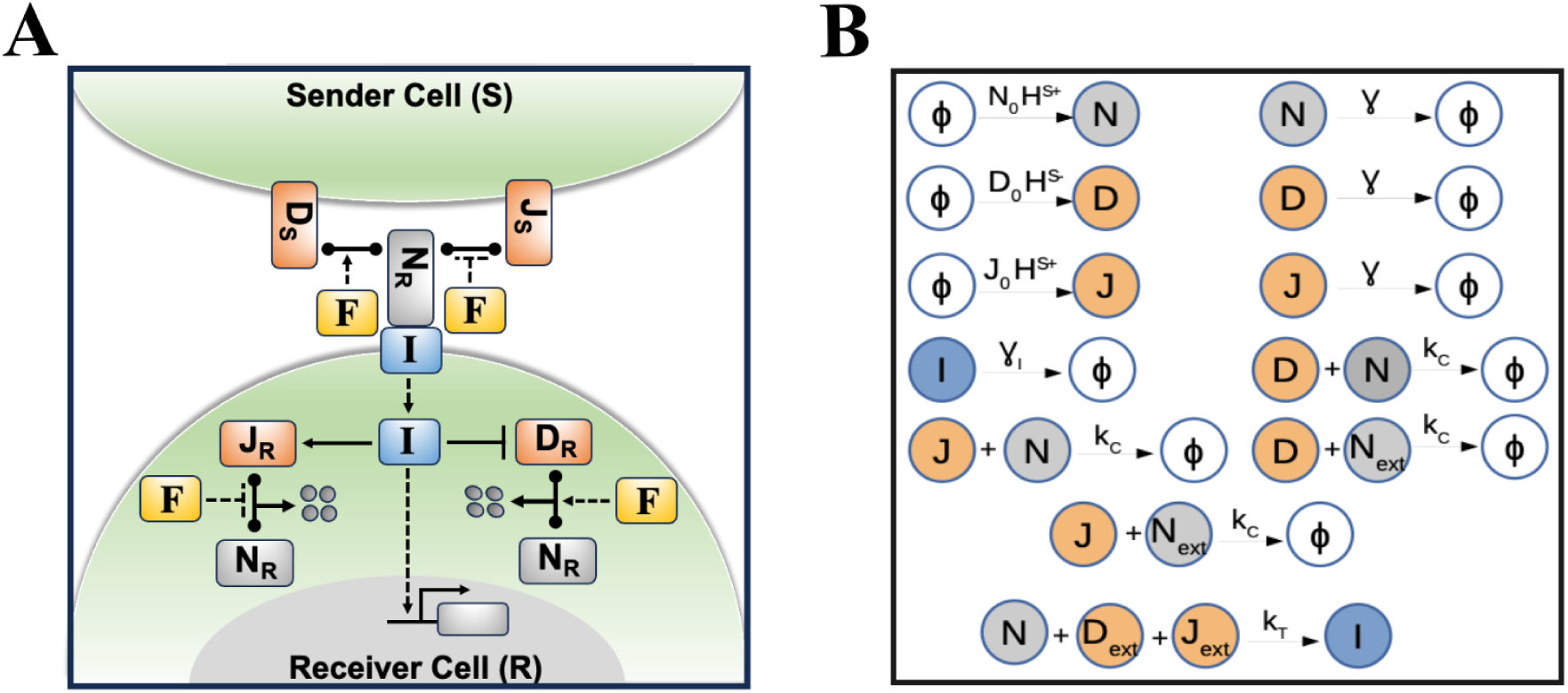
Detailed receptor-ligand (R/L) interactions between adjacent cells during Notch activation. A) An overview of the intra-cellular and inter-cellular interaction between a sender (S) and receiver (R) cell in the Notch signaling pathway. Ligands (Delta and Jagged) on the sender cell bind to Notch receptors on the receiver cell, triggering NICD production that regulates Notch, Delta, and Jagged under Fringe modulation. B) Reactions specifying the stochastic Notch-Delta-Jagged model (eq (1)), where *φ* is the empty state, *N_ext_*, *D_ext_*, and *J_ext_* represent the neighbouring cells’ Notch, Delta, and Jagged, respectively. *F* refers to the modulatory effect of Fringe on the receptor ligand binding affinity.

However, in the above equations, it is assumed that the Notch receptor has similar interaction strengths (both cis-inhibition (*k_C_*) and trans-activation (*k_T_*)) towards Delta and Jagged. Experimentally, it has been found that Notch can be glycosylated by Fringe (Morales et al 2002), which creates an asymmetry in the Notch binding affinity, thus defining two subpopulations of Notch: 1) Fringe-modulated Notch and 2) unmodulated Notch (LeBon et al 2014, Panin et al 1997). The glycosylated Notch has a higher binding affinity for Delta but a lower one for Jagged than the unmodified one, for both cis and trans interactions. In our augmented model, we will further assume that glycosylated Notch increases with NICD (*I*), as NICD activates Fringe. The corresponding deterministic equations (Boareto et al 2015) are now given by:

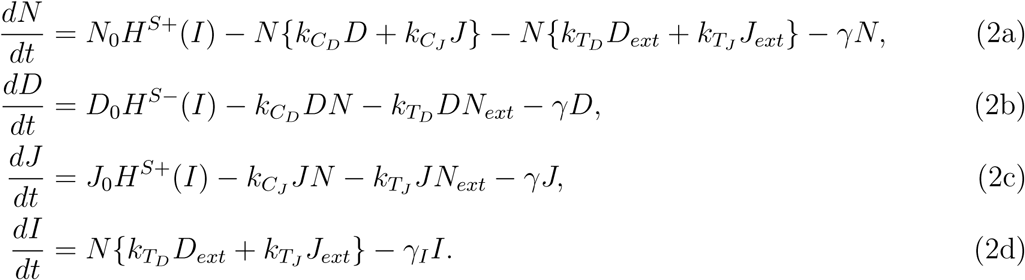

Here, the unequal cis-inhibition rates 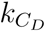 and 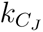 and trans-activation rates 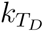 and 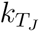 are functions of NICD (I); schematically, *k*(*I*) = *k*_0_[1 + *aH*^+^(*I*)] = *k*_0_*H^S^*^+^(*I, λ^F^*), *λ^F^* = 1 + *a*. The increased effect of Fringe with higher NICD is represented by a shifted Hill function *H^S^*^+^(*I, λ^F^*) and *λ^F^* represents the fold-change due to the Fringe activity for both tran-activation and cis-inibition; specifically, 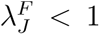 and 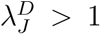 (Hicks et al 2000, Shimizu et al 2001). In the case of two interacting cells, the variables *N_ext_*, *D_ext_* and *J_ext_* need to be replaced by *N*, *D*, *J* of the neighboring cell. For more details of the model derivation (Lu et al 2013, Boareto et al 2015) see **SI Text S1.1.2**.

## 4 Stochastic Formulation

Most models for Notch-Delta signaling consider only the deterministic dynamics and the effects of extrinsic noise (Jolly et al 2015). Here, we consider intrinsic fluctuations due to the microscopic chemical processes contained in the system (1), as depicted by (Fig. 1B). The microscopic dynamics can be modeled as a four-dimensional Markov process in continuous time, which can be described by its master equation (Van Kampen 1992):

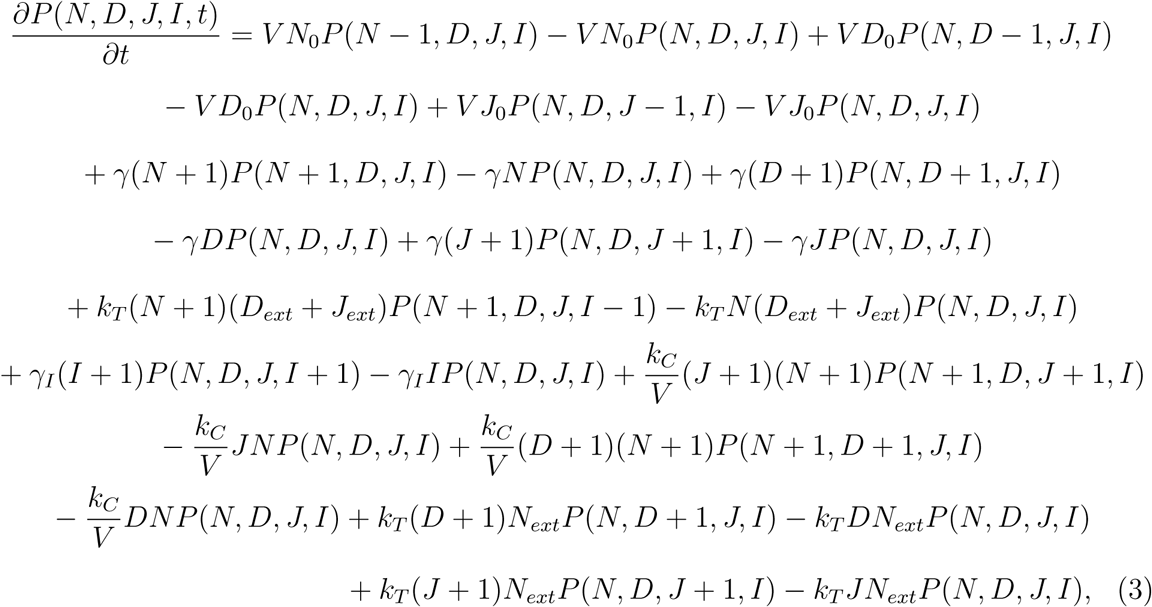

where *P* (*N, D, J, I, t*) is the probability of a particular state *N*, *D*, *J*, and *I* at time *t*, and *V* is the volume of the site in which the processes occur. This is given for the symmetric model, but has an obvious generalization to the non-symmetric case. A detailed derivation is provided in the **SI Text S2**.

## 5 Results

### 5.1 Early warning signals in forecasting transition for the Notch system

Initially, we ignore any possible role for Jagged. This leads to a simplified version of eq (1), given by:

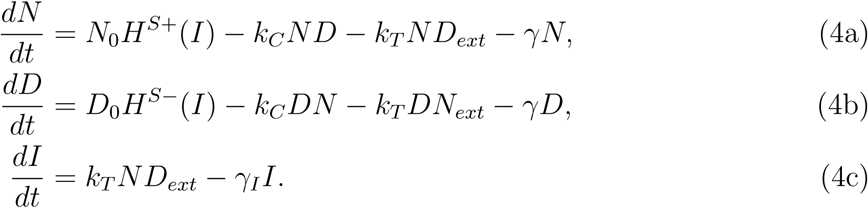

First, we performed a bifurcation analysis of the deterministic Notch-Delta circuit (eq 4) with varying external delta (*D_ext_*). The values of all model parameters of this circuit are presented in the **SI Table S6**. For small concentrations of *D_ext_*, the circuit exhibits a monostable state, termed a Sender (*S*), and for large values of *D_ext_*, the circuit exhibits another monostable state, termed a Receiver (*R*). Specifically, at high *D_ext_*, Notch signaling is activated, thus leading to higher levels of receptor Notch and lower levels of ligand Delta. Thus, this cell acts as a Receiver (R). Conversely, at low *D_ext_*, NICD levels are low, and consequently Notch levels are low, and Delta levels are high. Furthermore, for an intermediate concentration of (*D_ext_*), the circuit exhibits bistability between *S* and *R* states **(Fig. 2A)**, (Boareto et al 2015). This multi-stability arises through a saddle-node bifurcation and a hysteresis loop, which can be responsible for critical transitions in the presence of small stochastic perturbations.

**Figure 2.**
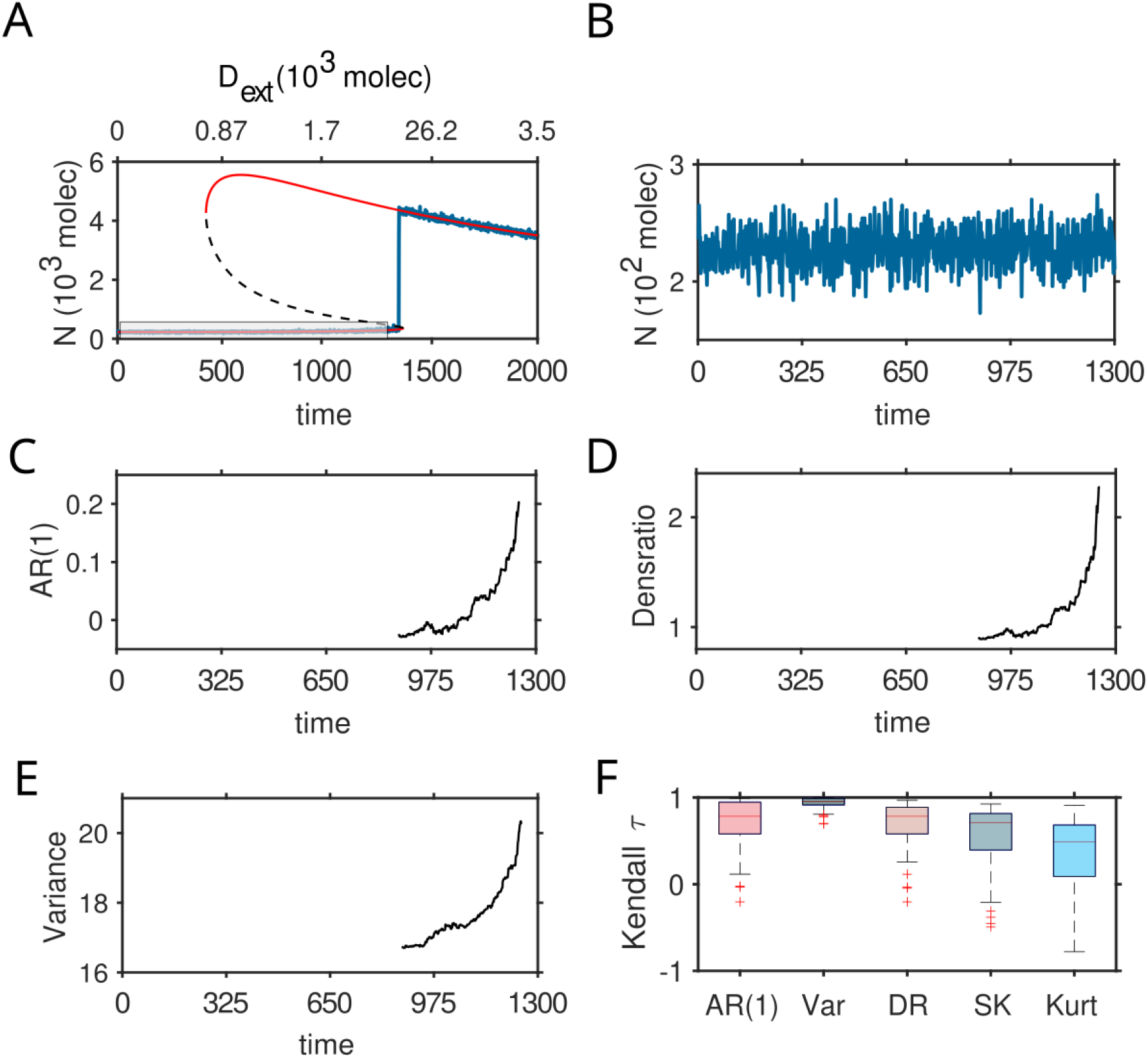
Detection of critical transitions between a low Notch level and a high Notch level in Notch-Delta circuit (eq (4)), using CSD-based indicators. (A) Transitions from Sender *S* state (low Notch, high Delta) to Receiver *R* state (high Notch, low Delta). Solid (red) lines indicate stable steady states, and dashed (black) lines indicate unstable steady states of the corresponding deterministic model. Stochastic time series are indicated by fluctuating lines. (B) Time series segment of the system before the transition to *R* state (represented by a rectangular box in (A). CSD-based indicators calculated from the time series after using a rolling window of 70% of data length: (C) AR(1), (D) Density ratio, and (E) Variance. (F) Kendall-*τ* rank correlations obtained from 100 time series segments of different CSD indicators, autocorrelation at lag-1 (AR(1)), variance (Var), density ratio (DR), skewness (SK), and Kurtosis (Kurt). The boxplots include the median, and the bottom and top edges of the box indicate the 25th and 75th percentiles, respectively. Box whiskers indicate the minimum and maximum values (see **SI Text S2 & S3**).

Next, in order to capture possible critical slowing down (CSD) in this circuit, we calculated the early warning signals (EWS) **(Fig. 2B)** of critical transitions in data sets obtained from stochastic simulations (see **SI Text S2.1.1 & Text S3.1**). The stochastic time series starting from a low Notch level (Sender state), with gradually increasing *D_ext_* (varying from 0 molecules to 3.5K molecules), exhibits a critical transition from Sender state *S* to Receiver state *R*. It should be mentioned in passing that the rate at which the *D_ext_*increases may have a significant impact on the nature of CSD indicators (see next section 5.2).

To determine several widely used early warning indicators including variance, autocorrelation at lag-1 (AR(1)) and density ratio (DR), we considered a time series segment (depicted in figure by a black box) before the transition to the receiver (*R*) state **(Fig. 2B)**. We calculated auto-correlation at lag-1 (AR(1)) **(Fig. 2C**), density ratio **(Fig. 2D**) and variance **(Fig. 2E**) using rolling windows 70% of the size of the time series (Dakos et al 2012) (see **SI Text S3.2**). Histogram plots show the distribution of trend statistics in the range *−*1 to 1. All early warning indicators show a signature of CSD trends. In particular, we found that all indicator values are increasing **(Fig. 2(C-E))**. An increase in EWS indicators is a well-known signature of an impending critical transition (Scheffer et al 2009, 2012). Furthermore, to assess the robustness of the EWS indicators, we generated 100 simulated independent stochastic trajectories and calculated the nonparametric trend statistic, Kendall’s *τ*, for AR(1), variance, density ratio, skewness, and kurtosis (Chen et al 2022). A box plot shows the distribution of trend statistics calculated from stochastic trajectories. The distribution of the rank correlations shows that the statistical trend for variance is stronger than other indicators, as Kendal’s *τ* values for the variance are clustered around 0.9 **(Fig. 2F)**. The trend statistics for all the indicators are positively distributed, and a positive statistical trend for an indicator shows an increasing trend in EWS of an upcoming critical transition (Sarkar et al 2021).

### 5.2 Robustness of CSD Indicators

Now, the system can evolve at any particular rate, which may influence the strength of these EWS indicators, depending on the inherent nature of the system. It has been evident that slowing down the rate of forcing may improve the performance of certain CSD indicators for systems exhibiting fold and transcritical bifurcations. (Clements and Ozgul 2016) Hence, selecting an appropriate rate for effective detection of CSD indicators is essential to identify transitions in the Notch system. However, quantifying such a rate is quite challenging for any complex biological process, as it depends on various factors, such as the ligand-binding mechanism, cooperative binding effect, degradation rates, and turnover rates (Sprinzak et al 2010, Nandagopal et al 2018, Gozlan and Sprinzak 2023). Here, we have investigated the robustness of CSD indicators by varying the forcing rate, which is depicted in **Fig. 3**. As expected, the analysis reveals that reducing the rate increases the robustness of the indicators. However, a balanced transition rate is crucial. If the transition occurs too slowly, it may prevent the proper functioning of an EWS indicator (Pavithran and Sujith 2021). Here, for this one-cell ND system, traditional EWSs perform well at forecasting an impending transition across a wide range of system response rates.

**Figure 3.**
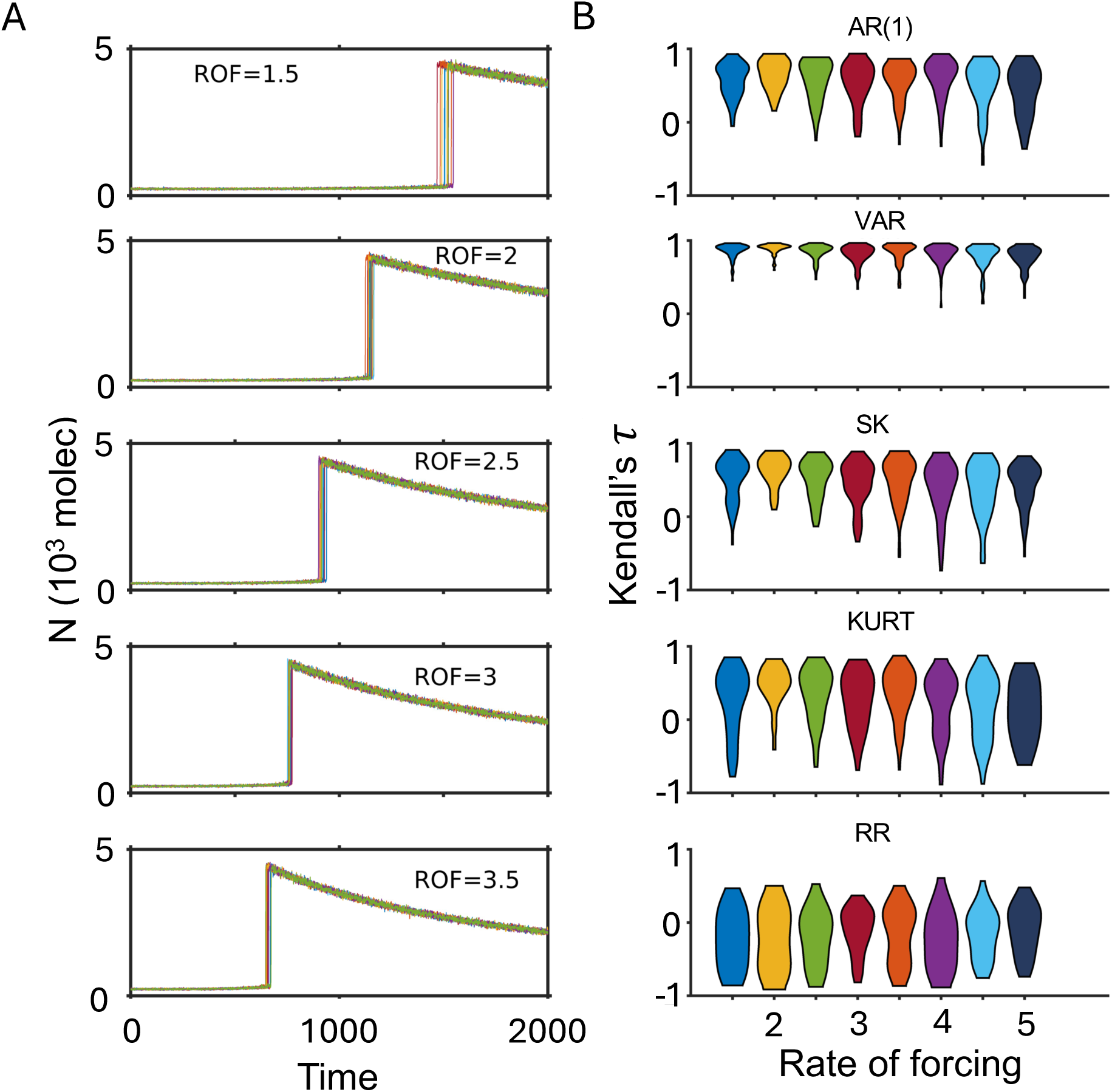
Robustness of the CSD-based indicators depend on the rate of forcing (ROF) for system (eq 4). (A) Transitions are observed from the Sender *S* state to the Receiver *R* state at different increasing rates of *D_ext_*(ROF). (B) Distribution of Kendall rank correlation values at different ROF. 100 independent stochastic trajectories are considered for each ROF (see **SI Text S3.2**).

### 5.3 Notch-Delta Signaling in a Two-Cell System

Next, we have investigated the dynamics of ND model for a two-cell system under the influence of intrinsic noise. Depending on the parameter values, the system exhibits a super-critical pitchfork bifurcation with the variation of the production rate of Delta *D*_0_ (**Text S1.2.1, Table S6, Fig. S1A**). Based on the level of NICD (I), we define these three stable states: a symmetric unpatterned state where both cells are hybrid (*S/R*_1_*, S/R*_2_), and two patterned states (*S*_1_*, R*_2_), (*R*_1_*, S*_2_). For increasing production rates (*D*_0_), the system first exhibits the monostable hybrid state (*S/R*_1_*, S/R*_2_); a further increase in *D*_0_ leads to bistability between Sender and Receiver states. In the presence of intrinsic stochastic fluctuations (**Text S2.2.1, Fig. S1A**), the system undergoes a second-order phase transition with the variation of *D*_0_.

To quantify critical slowing down (CSD) for such a second-order phase transition, we have calculated EWS (**Text S3.2**) from a trajectory obtained from a stochastic simulation of the two-cell system. The stochastic time series presents NICD levels with a continuously increasing production rate *D*_0_. We focused on the robustness of EWSs indicators to positively signal the impending transition from hybrid (*S/R*_1_*, S/R*_2_) state to either the Sender or the Receiver state, by measuring non-parametric correlations, using Kendall’s *τ*. For the analysis, we considered 100 independent stochastic times as previously and calculated Kendall’s *τ* for the indicators AR(1), variance (VAR), skewness (SK), kurtosis (KURT), return rate (RR), and density ratio (DR) using 70% rolling window sizes. Boxplot shows the distribution of Kendall’s *τ* rank correlation (**Fig. S1B**). We find positive rank correlations only for the variance, whereas other indicators exhibit both positive and negative correlations. These observations imply that a continuous transition from the symmetric state to a symmetry-broken state might not be strongly reflected in most CSD indicators.

## 6 Flickering - an alternative critical transition indicator for the Notch system

So far, we considered a single-ligand scenario, however, Delta and Jagged can have distinct production rates (Benedito et al 2009). We therefore incorporated NICD-mediated asymmetric regulation, wherein NICD activates Jagged and represses Delta, while assuming identical cis and trans interaction rates for Notch-Delta and Notch-Jagged interaction (Boareto et al 2015). Strong Jagged activation was implemented using a high Hill coefficient (*k_nj_ ≥* 5) (Boareto et al 2015). Under moderate to high *k_nj_* (*≥* 5), the model stabilizes a hybrid *S/R* state with significant existence of flickering (**Fig. S2(A-C)**), whereas reducing *k_nj_* abolishes the *S/R* state and eliminates flickering (**Fig. S2(D-F)**). Accordingly, *k_nj_* = 5 is used as the reference parameter set (**Table S6**) or WT for this study. The bifurcation analysis for the single-cell NDJ system (**Text S1.1.1**) depicts the appearance of a hybrid Sender/Receiver (S/R) state due to the asymmetric regulation, and thus this system acts as a three-way switch depicting three possible stable steady states: Sender (S), hybrid-Sender/Receiver (S/R), and Receiver (R) (Boareto et al 2015, Mukherjee and Levine 2023) **(Fig. 4A)**. When the stochastic trajectory of Notch was followed over a range of external Jagged *J_ext_* (eq (1)), instead of a smooth transition as was seen in the ND system, the stochastic trajectory **(Fig. 4A)** “flickers” between the Sender and hybrid *S/R* states (flickering is indicated in rectangular boxes, **Fig. 4B)**. Thus, flickering can act as an emerging alternative mechanism to critical slowing down in explaining rising trends in variance, lag-1 autocorrelation, and skewness (Wang et al 2012).

**Figure 4.**
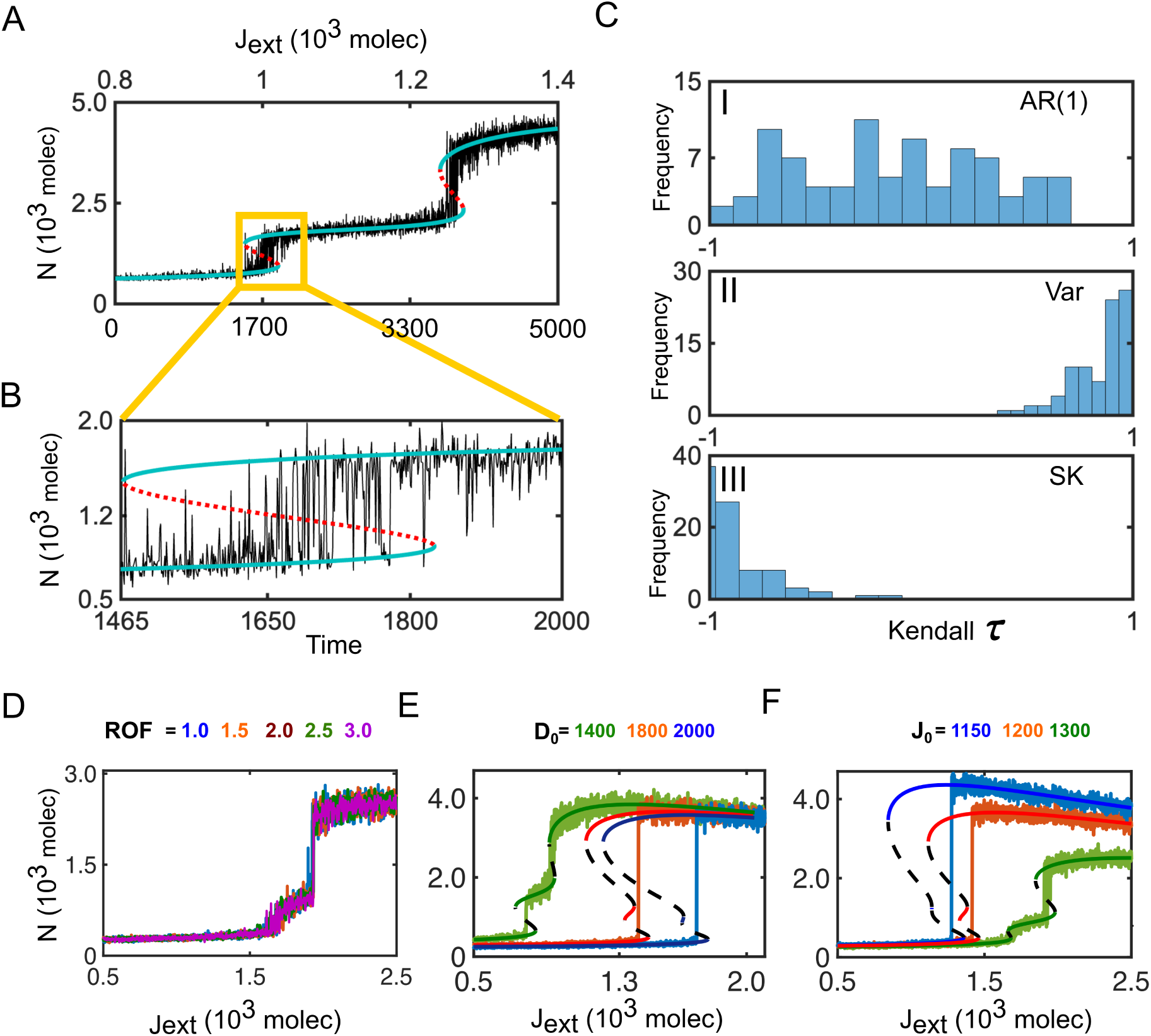
Detection of CSD-based indicator and flickering in Notch-Delta-Jagged signaling circuit (eq (1)). (A, B) Transitions from Sender S state to hybrid S/R state and S/R state to Receiver R state for a one-cell NDJ system (eq (1)). Solid (blue) lines indicate stable steady states, and dotted (red) lines indicate unstable steady states. Stochastic time series are indicated by fluctuating lines. (C) Kendall’s *τ* for (I) AR(1), (II) variance (VAR), and (III) skewness (SK), respectively, based on 100 replicates of trajectories. (D) Transitions are observed from the Sender *S* state to the Receiver *R* state at different increasing rates of *J_ext_* (ROF). Bifurcation diagram and stochastic time series observed in one-cell NDJ system in case of *n_j_* = 4 (Reference system, RS) by varying the (E) production rate of Jagged (*J*_0_ = 1150, 1200, 1300) and (F) Delta (*D*_0_ = 1400, 1800, 2000) individually (see **SI Text S2 & S3**).

We generated a set of independent stochastic trajectories (**Text S2.1.2**) in the region indicated by a box in **(Fig. 4(A-B))** and then used a Monte-Carlo simulation (**Text S3.1**) to calculate the trend statistics for AR(1), variance, and skewness. Trend statistics Kendall’s *τ* of the indicator AR(1), are evenly distributed between *−*1 and 1 **(Fig. 4CI)**, suggesting AR(1) is not a reliable estimator for flickering (Dakos et al 2013). The distributions of Kendall’s *τ* for variance (Var) and skewness (SK) are clustered around 1 and *−*1, respectively **(Fig. 4C(II, III))**. The positive trend of variance and the negative trend for skewness suggest that CSD-based indicators are not particularly effective in detecting transitions in this case. As discussed in the previous section, the property of CSD indicators may depend on the forcing rate for the ND circuit **(Fig. 3)**. Since our initial NDJ simulations were conducted with an arbitrary response rate, this rate may not be suitable for achieving good performance on the indicators. To explore this further, we examined transitions across a range of forcing rates. Surprisingly, neither the nature nor the timing of the transition was significantly affected by variations in this rate **(Fig. 4D)**. This may be attributed to flickering, which reflects the system’s inherent noise and stochasticity. In such a noisy model, the ensemble of stochastic trajectories carries a distinct signature; the traditional CSD indicators do not work well, and flickering becomes a significant precursor for capturing transitions. Hence, identifying the factors that regulate flickering is essential.

### 6.1 Ligand production rate controls flickering

Experimental studies have shown that ligand production rates play a critical role in shaping tissue-level patterning in the Notch signaling system (Boareto et al 2015, Mukherjee and Levine 2023, Bray 2006, Hunter et al 2016, Pan and Rubin 1997, Shaya and Sprinzak 2011). In particular, Boareto *et al*. demonstrated that a relative increase in Delta production promotes a *salt−and−pepper* pattern across tissues, while a higher production rate of Jagged suppresses this pattern and drives the cell to adopt similar *S/R* fates(Boareto et al 2015). This has been replicated in the one-cell model by simulating the intercellular interactions, considering the membrane-bound ligands of neighboring cells as external signals *D_ext_/J_ext_*.

We previously showed that a high Hill coefficient (*k_nj_* = 5, WT) induces persistent flickering (**Fig S2(A-C)**). To dissect the origin of flickering, we defined a near flicker free control state at *k_nj_* = 4, (**Fig S2(D-F)**) and used it as a reference system (RS-1) to systematically perturb model parameters and identify conditions that restore flickering and alter cell-fate decision dynamics. Within this framework, the NDJ system exhibits distinct dynamical behavior when simulated under varying ligand production rate (**Fig. S3**) (*D_ext_/J_ext_*). Interestingly, we found that it is possible to drive the baseline system (*k_nj_*= 4, RS) into a favorable *S/R* state by adjusting ligand levels, thereby supporting the re-emergence of flickering. Specifically, when the production rate of Delta is decreased, the enhanced noise in the system enables flickering but only from the *S/R* state to the receiver state (**Fig. 4E**, green color; **Fig. S3A**). Conversely, at relatively higher Delta production rates, the S/R state nearly disappears, substantially minimizing the ability of the system to flicker from the sender to receiver state (**Fig. 4E**, blue color; **Fig. S3A**). In contrast, at the relatively highest production rate of Jagged, the relative distance between the sender to receiver state decreases, enabling the system to again show a significant amount of flickering (**Fig. 4F**, green color; **Fig. S3B**). This result suggests that, in general, higher Jagged levels reshape the system’s dynamical landscape, leading to greater noise and variability in the Notch signaling pathway.

In summary, bifurcation analysis combined with stochastic simulations reveals that high Delta production suppresses flickering, driving stable commitment to either sender or receiver states, consistent with salt-and-pepper pattern formation at elevated Delta levels Boareto(Boareto et al 2015). In contrast, at relatively higher levels of Jagged production, the system displays extensive flickering, enabling it to fluctuate between the sender and the hybrid *S/R* state.

### 6.2 Stochastic Potential Analysis Indicate Strong Flickering Activity

Previous studies have shown that flickering arises when two alternative basins of attraction lie in close proximity in state space (Hicks et al 2000, Wang et al 2012, Scheffer et al 2009). However, stochastic time series alone often do not clearly reveal the relative proximity of these basins. To more directly characterize flickering and resolve the underlying state-space structure, in this section, we therefore performed stochastic potential analysis, which provides a clearer and more quantitative visualization of basin geometry.

The stability of different states of a complex dynamical system can be approximately represented by calculating an effective potential. The lowest value of the potential reflects the greater likelihood of the system existing in that particular state within a multi-stable region. When potential wells are shallow or closely spaced and combined with sufficient noise, the system can easily undergo stochastic switching between the nearby stable states creating flickering. Here, we have formulated the stochastic potential (Sarkar et al 2019) of *n_J_*= 4 (RS-1) at different *J_ext_*close to the transitions (**Fig. 5**). At high Delta levels, the potential landscape is characterized by deep, well-confined wells (**Fig. 5**A), indicating robust and stable attractor states. Decreasing Delta production destabilizes these wells (**Fig. 5B**, blue color; **Fig. S4(A,B)**) and at relatively lower Delta levels, the potential wells become stochastically distributed across hybrid S/R and receiver states (**Fig. 5**BIII), reflecting noise driven transitions. In comparison, increased Jagged production results in a more uniform and broadened distribution of potential wells, facilitating stochastic switching among sender, hybrid S/R, and receiver states (**Fig. 5C**, green color; **Fig. S4(C,D)**). Together, the stochastic potential landscape provides a unified, quantitative framework for linking noise-driven fluctuations to the geometry of underlying attractor basins based on ligands level, thereby revealing how flickering emerges from shallow or closely spaced potentials.

**Figure 5.**
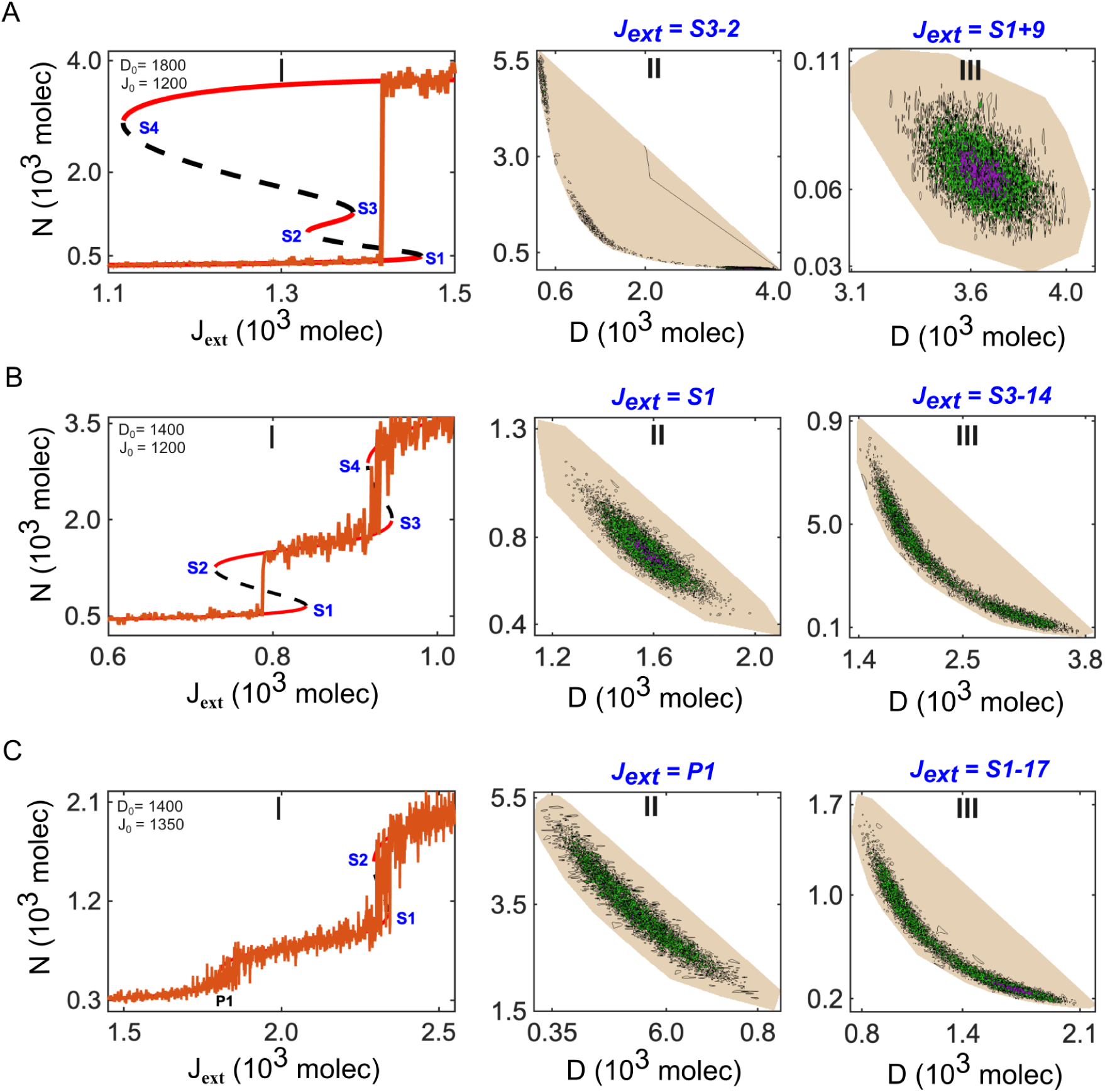
Potential landscape of the NDJ circuit (Eqn. (1)) under varying *J_ext_*. (A) High Delta production regime (*D*_0_ = 1800). (I) Bifurcation diagram showing the steady-state Notch level as a function of *J*_ext_. The deterministic system exhibits multiple saddle points (*S*1 = 840*, S*2 = 729*, S*3 = 944*, S*4 = 915), indicating transitions between alternative dynamical states. Solid red curves, dashed black curves denote denote stable and unstable equilibria. (II, III) Stochastic potential landscapes for representative values of *J*_ext_ corresponding to S1 (II) and S3 (III). Deeper colors (pink denotes the deepest potential, followed by green, black, and light gray) indicate deeper potential wells (local minima) representing stable attractors. (B) Low Delta production regime (*D*_0_ = 1400). (I) Bifurcation diagram of the steady-state Notch level versus *J*_ext_, showing a qualitatively altered bifurcation structure (*S*1 = 1461*, S*2 = 1329*, S*3 = 1382*, S*4 = 1116). (II, III) Corresponding stochastic potential landscapes for different values of *J*_ext_. (C) High Notch production regime (*J*_0_ = 1350). (I) Bifurcation diagram of the steady-state Notch level versus *J*_ext_ (*S*1 = 2337*, S*2 = 2292) and (II, III) corresponding stochastic potential for different values of *J*_ext_, (*P* 1 = 1820). Under a higher Jagged production rate, flickering mediated uniform distribution of stochastic potential landscapes. In (III), flickering generates multiple shallow local minima, reflecting stochastic switching among transient states (see **SI Text S3.3**).

### 6.3 Limitations of CSD Indicators in Two-cell Notch-Delta-Jagged Signaling

Tissue-level patterning is fundamentally a result of cell-cell communication. Here, we have next investigated the dynamical properties of NDJ signaling for two interacting cells (**Text S1.2.2**) as a function of the production rate of Jagged (*J*_0_) under different constant Delta production rates (**Fig. 6A**). At a lower *D*_0_ value (400, 600), the system exhibits monostability with a single stable branch. As *D*_0_ increases (800, 1400), the central branch splits into two unstable and one stable one, indicating a bistable regime characteristic of a super-critical pitchfork bifurcation (**Fig. 6A**). The NDJ system again exhibits a transition between symmetric hybrid state (*S*_1_*/R*_1_*, S*_2_*/R*_2_) and one of the patterned states; this is clearly visible through the time series obtained from the stochastic simulation (**Fig. 6B, Text S2.2.2**). Parenthetically, flickering is not seen in the two-cell model, as there is no range of stable state coexistence for a supercritical bifurcation. Similar to our study of the two-cell Notch-Delta case, we investigated the phase transition by calculating the rank correlations of different CSD-based EWS indicators for NDJ signaling. The rank correlation of Kendall *τ* is obtained from a stochastic time series representing NICD levels with a continuously decreasing production rate of Jagged. For the EWS analysis, we consider 100 independent stochastic time series segments before the transition from the (*S*_1_*/R*_1_*, S*_2_*/R*_2_) state to one of the asymmetric states. We also calculated similar rank correlations for the indicators AR(1), VAR, SK, KURT, RR, and DR using 70% rolling window size. For all indicators, we found both positive and negative correlations obtained from ensembles of trajectories (**Fig. 6C**). This again indicates that CSD-based early warning signals are not effective in predicting abrupt transitions associated with supercritical pitchfork bifurcations. Therefore, in the later section, we focus exclusively in the NDJ system on the single-cell case only.

**Figure 6.**
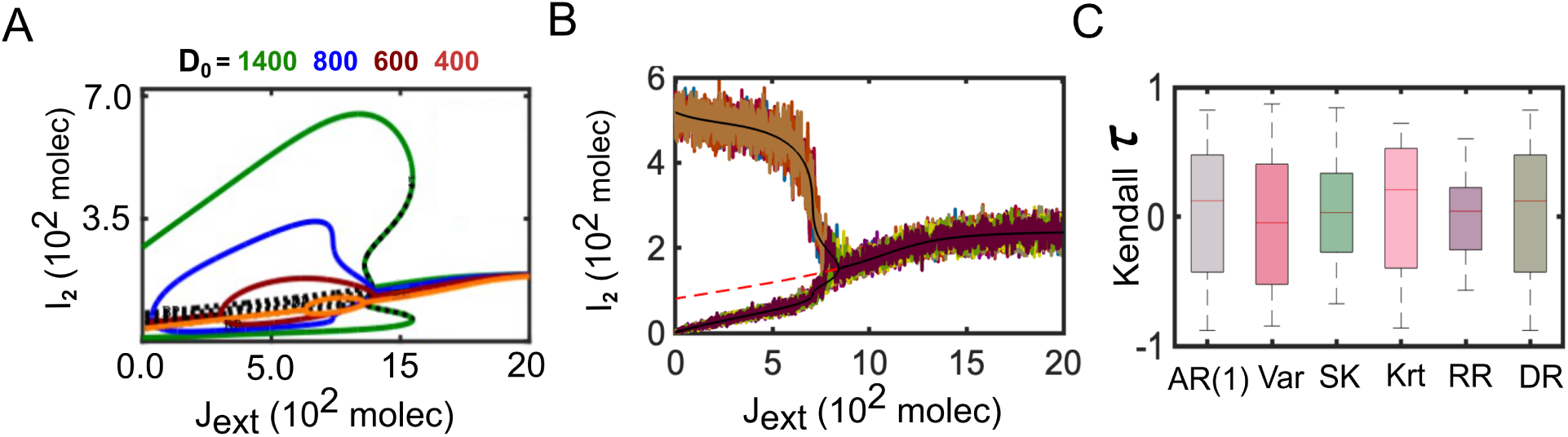
Bifurcation Diagram and transition between different cell states for two interacting NDJ cells. (A) Changes in the topology of the bifurcation diagram, while varying the production rate of Delta (*D*_0_ = 400, 600, 800, 1400). The colored solid lines indicate stable steady states, and the black dotted lines represent unstable steady states. (B) Bifurcation diagram and stochatic simulation at *D*_0_ = 1000. The solid black line, dotted red lines, and fluctuating lines represent stable states, unstable states, and 100 stochastic time series, respectively. (C) Kendall’s *τ* rank correlations of autocorrelation at lag-1 (AR(1)), variance (Var), skewness (SK), kurtosis (Kurt), return rate (RR), and density ratio (DR). The box plots display the median, with the bottom and top edges of the box representing the 25th and 75th percentiles, respectively (see **SI Text S2 & S3**).

### 6.4 Modulation of Flickering via Fringe-mediated Asymmetric binding

In this section, we investigate how the Fringe-mediated asymmetry in ligand-Notch binding affinity contols flickering and, ultimately the cell fate choices. It is well established that Fringe-mediated glycosylated Notch enhances its binding affinity for Delta, while reducing its affinity for Jagged, an effect that is incorporated into our model as 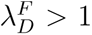 and 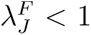 (**SI Text S1.1.2**) (Hicks et al 2000, Shimizu et al 2001, Mukherjee and Levine 2023). By varying these two parameters (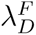 and 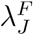), we simulate the binding preferences of distinct Notch variants modulated by different variants of Fringe (Boareto et al 2015). First, in the absence of Fringe activity (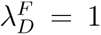 and 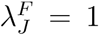), the starting system exhibits three possible stable steady states: Sender (S), hybrid-Sender/Receiver (S/R) and Receiver (R) (Boareto et al 2015, Mukherjee and Levine 2023) with respect to varying external Delta (*D_ext_*) (**Fig. 7A**). Under these conditions, the stochastic trajectory of Notch displays strong flickering behavior, which is further supported by uniform distributions in the stochastic potential landscape (**Fig. 7(A,B), Fig. S5A, Fig. S6A**). Moreover, trend statistics using Kendall’s *τ* for critical slowing down (CSD) indicators - AR(1), variance, and skewness show patterns similar to those observed for the NDJ case (**Fig. 7C**).

**Figure 7.**
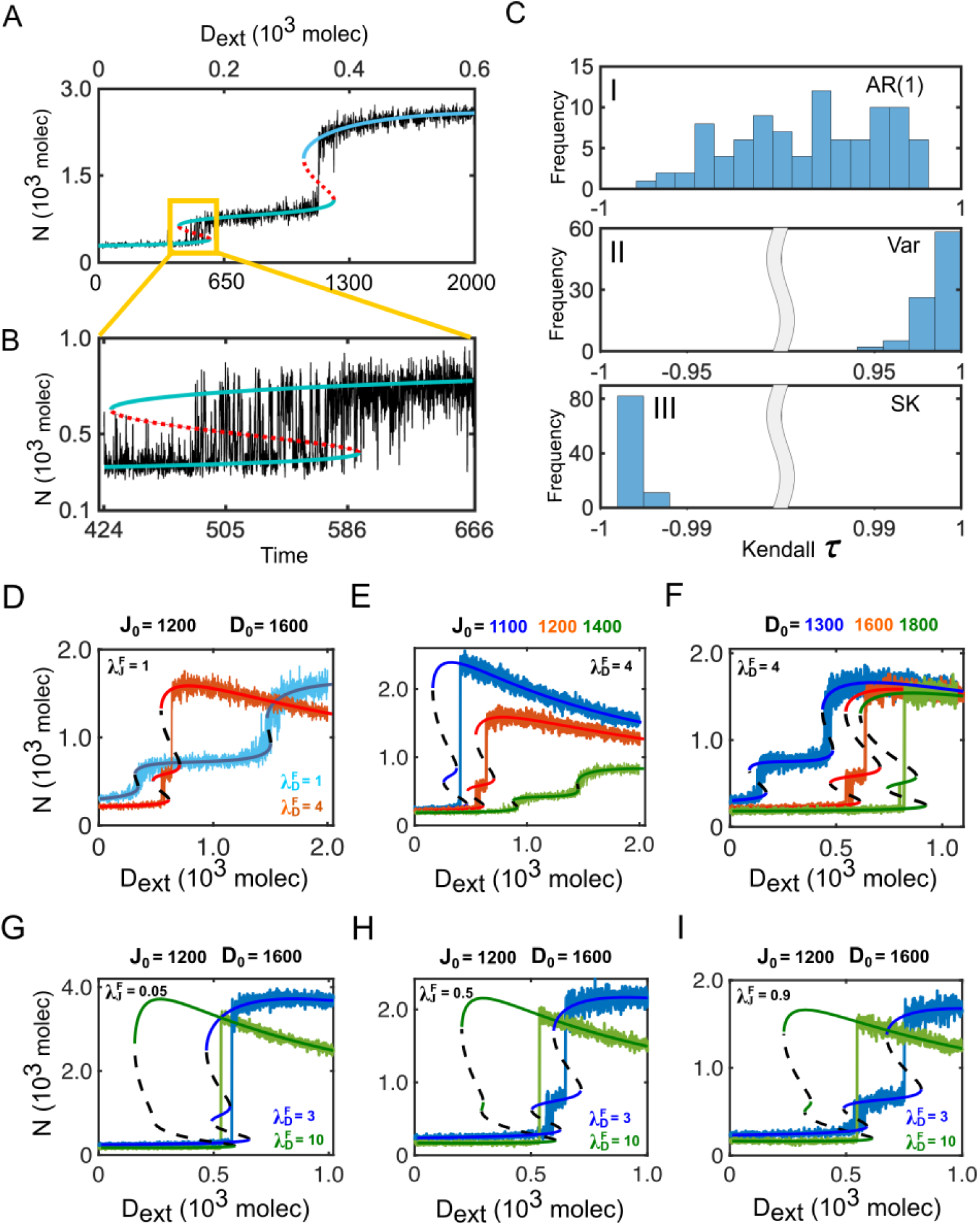
Detection of critical transition in Fringe mediated NDJ system (eq 2) and controllng of flickering via altering ligand production rate in the presence and absence of Fringe effect. (A) Transitions from Sender S state to hybrid S/R state and S/R state to Receiver R state for a one-cell NDJ system (eq 2). Solid blue lines, dotted red lines and fluctuating lines indicate stable steady states, unstable steady states and stochastic time series respectively. (B) A segment in A, near saddle-node bifurcation is indicated by the boxed region. (C) Performance of CSD indicators of 100 replicates of trajectories. Kendall’s *τ* for (I) AR(1), (II) variance (VAR), and (III) skewness (SK), respectively. (D,E,F) Bifurcation diagram and stochastic time series for the NDJ system (*n_j_* = 5, WT) under varied Fringe effect on only Delta binding (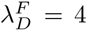 and 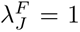). Under this condition, quantification of flickering under the varied production rate of (E) Jagged and (F) Delta. (G,H,I) Bifurcation diagram and stochastic time series for the NDJ system (*n_j_* = 5, WT) by varying Fringe effect on both Delta and Jagged binding under low 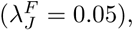 moderate 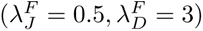 and strong effect 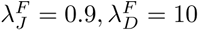) (see **SI Text S2 & S3**).

Next, a systematic analysis was conducted to investigate the effect of Fringe on Flickering dynamics. We first examined the effect of Fringe on Delta and Jagged individually (Case-1 and Case-2), and then explored the combined influence of Fringe on both ligands (Case-3). In the first scenario, where the Fringe influences Delta binding but not Jagged binding (Case-1, 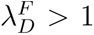 and 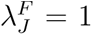), as the binding affinity between Notch and Delta increases 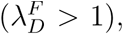 the system exhibits notable alteration in dynamical behavior, reducing flickering (**Fig. S(5,6)**). Specifically, at 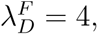 flickering is nearly eliminated (**Fig. 7D**, magenta color; **Fig. S5D**). Furthermore, the stochastic potential analysis illustrates this clearly, where the initial uniform distributions become increasingly localized as 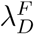 rises (**Fig. S6D**). This raises an important question: under what conditions, if any, can flickering re-emerge in this system? As previously observed in RS-1 (Section 6.1), altering the ligand production rate can restore flickering. Similarly, for Case-1, we found that relatively higher production rate of Jagged (**Fig. 7E**, **Fig. S7A**) and reduced Delta level enhances the system’s ability to transition back and forth between Sender to S/R to Receiver state, resulting in significant flickering (**Fig. 7F**, **Fig. S7B**). This can be predominantly observed from stochastic potential analysis (**Fig. S(8,9)**). On the other hand, for Case 2, we consider Fringe modulates only Notch-Jagged binding, not for Delta (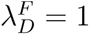 and 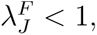 Case-2). Here, increasing Fringe-mediated inhibition on Jagged has a negligible impact on the system’s dynamics (**Fig. S10**).

Finally, we have investigated the system dynamics and stochastic behavior considering Fringe effects on both ligands binding (Case-3). When Fringe moderately impacts Delta binding to Notch 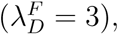 the occurrence of flickering depends on the threshold of Fringe’s modulation over Jagged (**Fig. 7(G-I)**). Under these parametric conditions, if Jagged binding is subject to low Fringe effect 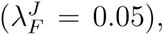 the system effectively behaves like a two-way switch (**Fig. 7G, blue color**). Flickering appears only when both Delta and Jagged binding are moderately influenced by Fringe, indicating a delicate balance required for dynamic switching (**Fig. 6(H-I)**, blue color; **Fig. S11(A,C)**). In contrast, with high Fringe impact on Delta binding 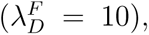 the system remains limited to sender and receiver states, no matter how Jagged binding is altered (**Fig. 7(G-I)**, green color; **Fig. S11(B,D)**). However, the Notch signaling response is also shaped not only through Fringe activity, but it also critically depends on the nature of signal transmission from the neighboring cells (*D_ext_/J_ext_*).

We have also investigated the dynamical properties of Fringe-mediated NDJ signaling for two interacting cells as a function of *J_ext_* under varying Delta production rates (*D*_0_) (**Fig. S12A**). With increasing *D*_0_, the bifurcation diagram transitions to a supercritical pitchfork bifurcation (**Fig. S12A**), like observed in **Fig. 6A**. Altering the Fringe-mediated effects on either Delta or Jagged binding modifies only the structure of the branches in the bifurcation diagram (**Fig. S12(B,C)**), without changing its overall qualitative nature.

### 6.5 Fine-tuning of Flickering by External Ligands Signal’s Strength

Since the Notch signaling pathway is primarily governed by external ligand signals from neighboring cells, this section focuses on characterizing its behavior under varying external inputs, both in the presence and absence of Fringe. When Fringe strongly regulates both ligands 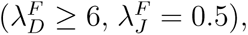 the system tends to remain stable in either the sender or receiver state (**Fig. 8(A,D)**, green color; **Fig. S13(A,B), Fig. S14(A,B)**). Under these conditions, adjusting the frequency of external signals fails to reinstate flickering (*D_ext_/J_ext_*). However, with moderate Fringe modulation 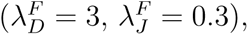 flickering can be adjusted by varying the inputs *D_ext_/J_ext_*. In this case, where Notch signaling is predominantly driven by Jagged (*J_ext_ ≥* 3000, **Fig. 8B**, **Fig. S13C**) rather than Delta (*D_ext_ ≤* 400, **Fig. 8E**, **Fig. S14C**), the system stabilizes in the hybrid *S/R* state, showing strong presence of flickering and allowing the cell to act as both sender and receiver. In contrast, when Fringe is absent (System-1), the system exhibits pronounced flickering even at relatively low levels of external Jagged or Delta (**Fig. 8(C,F)**, **Fig. S13D, Fig. S14D**). Also, it is noticeable that, under this case (system-1), a higher level of external jagged (*J_ext_* = 4000), drives the system toward a purely receiver state (**Fig. 8C**, **Fig. S13D**). Whereas, decreased external Delta levels (*D_ext_ ≤* 600) favor the sender state, while supporting a moderate coexistence of the *S/R* states (**Fig. 8F**, **Fig. S14D**).

**Figure 8.**
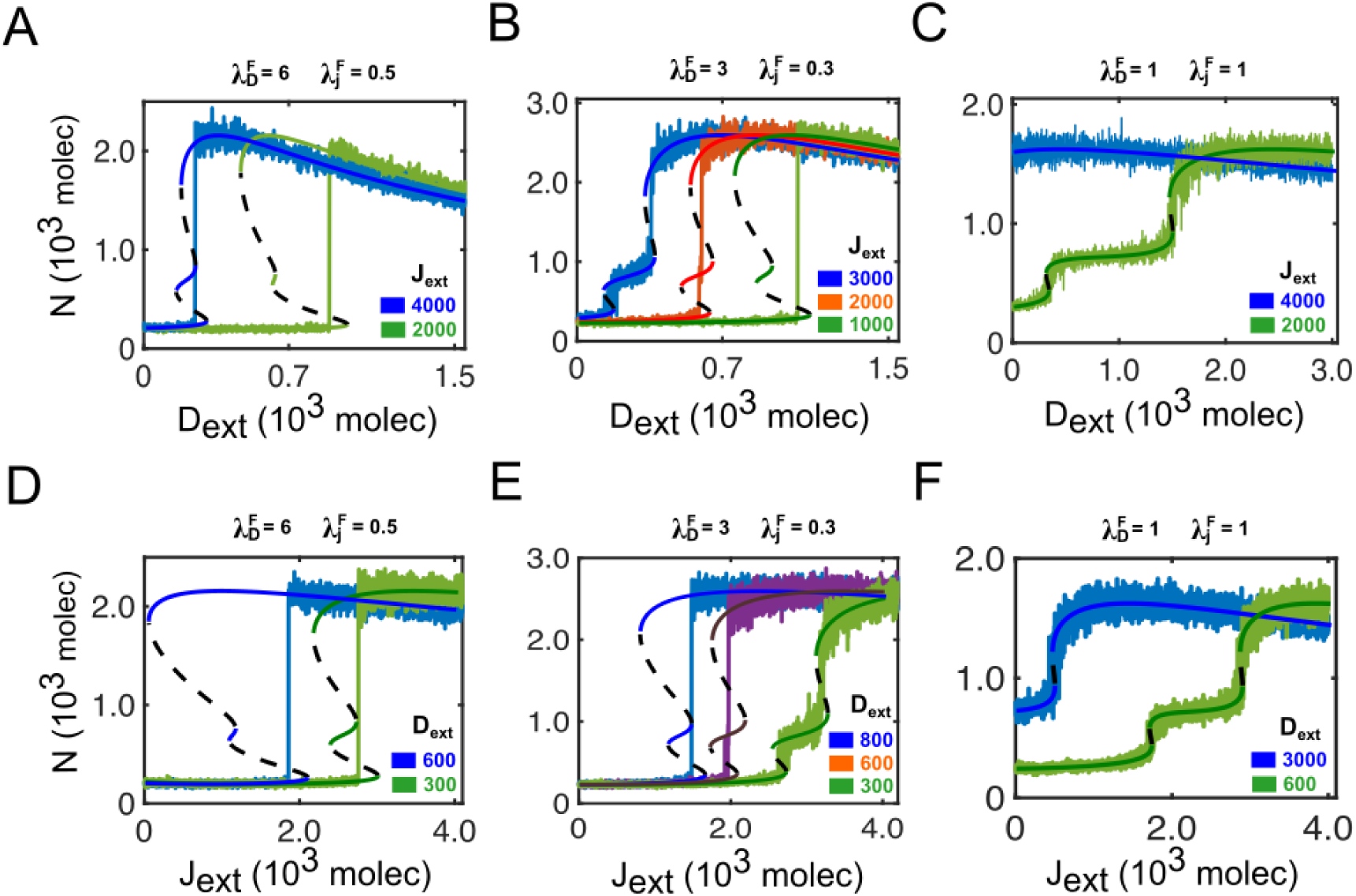
External signals modulate flickering of Fringe-mediated NDJ system (eq 2). Under the (A,D) strong 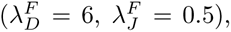 (B,E) intermediate 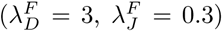 and (C,F) without Fringe 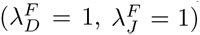 effect of Fringe on both delta and jagged, by varying external jagged (*A, B, C*) and external delta (*D, E, F*). The solid lines, dotted lines and fluctuating lines indicate stable steady states, unstable steady state and stochastic time series (see **SI Text S3.1**).

In summary, the persistence of the *S/R* state is shaped not only by NICD-driven asymmetry on Jagged but also by asymmetries caused by Fringe and external signals. Our analysis reveals that all these factors influence noise levels and, hence, flickering, increasing the likelihood of the system occupying multiple states simultaneously under certain conditions. This may bring challenges in developing rational drug design and create significant obstacles for targeted therapies.

## 7 Discussion

Sudden regime shifts are ubiquitous in complex systems; understanding their origins and forecasting them can be of paramount importance (Moris et al 2016, Shin and Cho 2023, Sarkar et al 2019, Scheffer et al 2012, Batt et al 2013). Such abrupt shifts can significantly impact systems in various ways. While studying the relationship between Waddington’s epigenetic landscape metaphor and genetic programmes in the context of cell fate decisions, Moris et al (2016) proposed an alternative view based on single-cell technologies (Pina et al 2012) that the actual change in cell fate occurs as a discontinuous transition; a sudden, discrete “jump” between stable states rather than a gradual slide. In recent years, predicting discontinuous transitions across various systems using early warning signals derived from measurement data has attracted significant attention within the scientific community. Prior studies on sudden transitions and their early warning indicators have considered demographic stochasticity (Sarkar et al 2019, Wang et al 2012). It is known that fluctuations in the abundance of molecules in living cells can play a fundamental role in the behavior of complex systems (Constable et al 2016). Therefore, it is important to move beyond deterministic approaches and consider the impact of intrinsic noise on the dynamics.

In this work, we systematically investigated how sudden transitions in Notch signaling dynamics can be anticipated under intrinsic noise and how the nature of these transitions depends on gene regulatory networks, ligand dynamics, and asymmetry. By combining deterministic bifurcation analysis with stochastic simulations and early warning signals, our analysis highlights the limitations of conventional CSD-based indicators. Our results uncover flickering as a dominant precursor of biological transitions in NDJ systems.

Further, our analysis found that in the ND circuit, transitions between sender (S) and receiver (R) states arise via saddle-node bifurcations and hysteresis. We find that early warning indicators, such as autocorrelation, variance, return rate, skewness, and kurtosis, can capture CSD during transitions among different phenotypes. These results align with theoretical expectations for fold bifurcations and suggest that cell-fate switching driven by external Delta can be predicted from noisy time-series data.

However, this is not universal, as in two-cell models of Notch signaling, transitions occur via a supercritical pitchfork bifurcation, and CSD indicators fail to show consistent trends. For a two-cell ND system, we find that only variance can capture the transitions. This observation reinforces the idea that continuous symmetry-breaking transitions do not necessarily exhibit strong critical slowing down and may therefore not be detectable using standard EWS frameworks. From a biological perspective, this implies that certain developmental patterning transitions, particularly those involving gradual symmetry breaking, may be inherently difficult to predict solely from fluctuations. Lastly, we have analyzed the effect of the rate of forcing on the strength of EWS. We have found that increasing the rate of force weakens the strength of indicators.

A central finding of our study is that including Jagged qualitatively alters the stochastic dynamics of the Notch circuit, consistent with other recent observations (Chen et al 2025). In the one-cell NDJ system, asymmetric regulation introduces a third hybrid *S/R* state, effectively transforming the circuit into a three-way switch. In this multistable regime, stochastic trajectories frequently hop between neighboring attractors before a final transition occurs. Occurrence of flickering is observed between the sender *S* state and the hybrid *S/R* state to the receiver *R* state. Thus, flickering emerges as a robust precursor of state changes, even when classical CSD indicators perform poorly or exhibit ambiguous trends.

Importantly, flickering is not simply a byproduct of noise intensity or forcing rate. Instead, it reflects the underlying geometry of the dynamical landscape; shallow and closely spaced potential wells allow noise-driven switching between states. Our stochastic potential analysis makes this explicit, showing how flickering corresponds to flattened or fragmented potential landscapes in which multiple attractors coexist with comparable stability. This interpretation is consistent with the attractor-based view of cell fate decisions, where progenitor and differentiated states correspond to stable or metastable attractors in a high-dimensional gene regulatory landscape (Jia et al 2017). Noise-driven excursions can transiently bring cells close to basin boundaries, and differentiation signals may act by flattening or destabilizing attractor wells, thereby facilitating stochastic jumps into neighboring lineage-specific attractors rather than enforcing deterministic trajectories. (Mojtahedi et al 2016, Galbraith et al 2022)

Our results further demonstrate that the relative production rates of Delta and Jagged act as key control parameters for flickering. High Delta production stabilizes sender-receiver patterning by enforcing cis-inactivation and lateral inhibition (Sprinzak et al 2011) and suppresses flickering, consistent with robust salt-and-pepper outcomes. This Delta-mediated lateral inhibition has been found to sharpen the Wnt7/*β*-catenin gradient into a stable boundary (Li et al 2013), enabling robust patterning. In contrast, higher Jagged production promotes a hybrid S/R state and enhances flickering, enabling frequent stochastic switching among the states. These findings suggest that the presence of the ligand can fundamentally reshape the noise-driven dynamics of the system.

Our analysis further revealed Fringe as a tunable regulator of robustness versus plasticity in Notch signaling. Fringe enhancement of Delta binding suppresses flickering and effectively reduces the NDJ circuit to a two-state switch. Moderate Fringe effects, however, preserve multistability and allow flickering to persist, particularly when combined with appropriate ligand production rates or external signals. By controlling binding asymmetries, Fringe can either stabilize fate decisions or permit noisy exploration of alternative states. This dual role may help explain how the same core Notch pathway supports both reliable pattern formation and flexible cell fate responses across different developmental and pathological contexts.

In summary, our findings demonstrate the impact of microscopic processes on the intercellular Notch signaling pathway. While CSD-based indicators are effective in certain regimes, they are insufficient for capturing the full range of behaviors in multistable, noise-driven systems such as NDJ signaling. Flickering, by contrast, emerges as a powerful and intuitive indicator of proximity to transitions in such contexts. In systems exhibiting flickering, increased variability may not signal a tipping point, but rather an intrinsically fluctuating landscape shaped by ligand dynamics, regulatory feedback, and post-translational modification. Accounting for these factors will be essential for rational intervention strategies aimed at stabilizing or redirecting cell fate decisions, where single-cell approaches are being investigated in detail (Islam and Bhattacharya 2025). More broadly, the framework developed here, combining the stochastic simulations, EWS analysis, and potential landscape reconstruction, can be extended to other intercellular signaling pathways and multicellular decision-making processes. Such approaches may prove valuable in the near future for understanding phenotypic heterogeneity in development, tissue homeostasis, and disease progression, as well as for synthetic approaches to multicellularity (Soĺe et al 2024).

## Acknowledgements

PSD acknowledges financial support from the Science & Engineering Research Board (SERB), Govt. of India [Grant No.: CRG/2022/002788]. HL acknowledges support from the NSF Center for Theoretical Biological Physics, grant NSF-PHY2019745. MKJ acknowledges support from the Ramanujan Fellowship, SERB, Govt. of India (SB/S2/RJN-049/2018), and by Param Hansa Philanthropies. BT was supported by the Institute of Eminence (IoE) post-doctoral fellowship awarded by the Indian Institute of Science (IISc).

## Conflict of interest statement

The authors declare no competing financial or non-financial interests.

## Notes

### Competing Interest Statement

The authors have declared no competing interest.

