## Supplementary Methods and Figures for "Early Detection of Sudden Transitions in Notch signaling"

### Supporting Information

**S4 Supplementary Figures** **19**

### S1 Text S1. Deterministic Models:

#### *S1.1 Text S1.1 One-Cell Model Equations:*

##### S1.1.1 Text S1.1.1 Notch-Delta-Jagged Circuit:

The Notch-Delta pathway is an evolutionarily conserved mechanism that governs cell fate during development. It is activated when the Notch receptor of one cell interacts with the ligand Delta or Jagged of its neighboring cell, which releases NICD. When Notch receptor ($N$) belonging to one cell can interact with both the ligands of the same cell ($D$ or $J$)- known as cis-inhibition and with those of the neighboring cell- Jagged or Delta- known as trans-activation. Additionally, Jagged is also included in the signaling circuit, with feedback effects mediated by NICD. The deterministic equations Boareto et al (2015) for the dynamics of Notch ($N$), Delta ($D$), Jagged ($J$), and NICD ($I$) are given by:

$$\frac{dN}{dt} = N_0 H^{S+}(I) - k_C N(D+J) - k_T N(D_{ext} + J_{ext}) - \gamma N, \tag{1a}$$
$$\frac{dD}{dt} = D_0 H^{S-}(I) - k_C DN - k_T D N_{ext} - \gamma D, \tag{1b}$$
$$\frac{dJ}{dt} = J_0 H^{S+}(I) - k_C JN - k_T J N_{ext} - \gamma J, \tag{1c}$$
$$\frac{dI}{dt} = k_T N(D_{ext} + J_{ext}) - \gamma_I I, \tag{1d}$$

where $N, D, J$, and $I$ represent Notch, Delta, Jagged, and NICD, respectively. $\gamma$ represents the degradation rate of all three proteins, Notch, Delta, and Jagged, and $\gamma_I$ the degradation rate of NICD. $k_C$ and $k_T$ are the strengths of cis-inhibition and trans-activation, respectively. $N_0$, $D_0$, and $J_0$ are the production rates of Notch, Delta, and Jagged, respectively. $N_{ext}$, $D_{ext}$, and $J_{ext}$ represent the amount of protein available for binding, which can be present on the membrane surface of neighbouring cells. Here, shifted Hill functions are used to represent the effect of NICD ($I$) on the production rates of the proteins. Shifted Hill functions are defined as $H^S(I, \lambda) = H^-(I) + \lambda H^+(I)$, where $\lambda$ represents the fold change in the production rate. If $\lambda > 1$ (for activation), then it is denoted as $H^{S+}(I)$ and if $\lambda < 1$ (for repression), then it is denoted as $H^{S-}(I)$. The details of the model can be found elsewhere (Boareto et al 2015, Sprinzak et al 2010). We performed the bifurcation analysis of this circuit for a fixed level of external Delta, $D_{ext}$. The parameters used for this analysis are given in the Table.

##### S1.1.2 Text S1.1.2 Fringe mediated Notch-Delta-Jagged Circuit:

The asymmetric Notch-Ligand binding is incorporated by the effect of Finge, thus the resulting model for one cell is given by :

$$\frac{dN}{dt} = N_0 H^{S+}(I, \lambda_{I,N}) - N\{k_{C_D} D + k_{C_J} J\} - N\{k_{T_D} D_{ext} + k_{T_J} J_{ext}\} - \gamma N, \tag{2a}$$
$$\frac{dD}{dt} = D_0 H^{S-}(I, \lambda_{I,D}) - k_{C_D} DN - k_{T_D} DN_{ext} - \gamma D, \tag{2b}$$
$$\frac{dJ}{dt} = J_0 H^{S+}(I, \lambda_{I,J}) - k_{C_J} JN - k_{T_J} JN_{ext} - \gamma J, \tag{2c}$$
$$\frac{dI}{dt} = N\{k_{T_D} D_{ext} + k_{T_J} J_{ext}\} - \gamma_I I. \tag{2d}$$

Where $k_{C_D} = k_C H^{S+}(I, \lambda_{F,D})$ and $k_{T_D} = k_T H^{S+}(I, \lambda_{F,D})$ with $\lambda_{F,D} > 1$.

##### S1.1.3 Text S1.1.3 Notch-Delta Circuit:

In the absence of Jagged in the above system reduces to a Notch-Delta circuit, and the corresponding deterministic model is as follows:

$$\frac{dN}{dt} = N_0 H^{S+}(I) - k_C ND - k_T ND_{ext} - \gamma N, \tag{3a}$$
$$\frac{dD}{dt} = D_0 H^{S-}(I) - k_C DN - k_T DN_{ext} - \gamma D, \tag{3b}$$
$$\frac{dI}{dt} = k_T ND_{ext} - \gamma_I I, \tag{3c}$$

##### S1.2 ***Text S1.2 Two-Cell Model Equations:***

In the next subsections, we describe the model for two interaction cells for the two models: Notch-Delta only (N-D), Notch-Delta-Jagged (N-D-J), and their corresponding stochastic systems.

###### S1.2.1 Text S1.2.1 Notch-Delta Module:

The dynamics of interaction between two cells Boareto et al (2015) are modelled as:

$$\frac{dN_1}{dt} = N_0 H^{S+}(I_1, \lambda_{I_1,N_1}) - N_1\left[(k_C D_1 + k_T D_2)\right] - \gamma N_1, \tag{4a}$$
$$\frac{dD_1}{dt} = D_0 H^{S-}(I_1, \lambda_{I_1,D_1}) - k_C D_1 N_1 - k_T N_2 D_1 - \gamma D_1, \tag{4b}$$
$$\frac{dI_1}{dt} = k_T N_1 D_2 - \gamma_I I_1, \tag{4c}$$
$$\frac{dN_2}{dt} = N_0 H^{S+}(I_2, \lambda_{I_2,N_2}) - N_2\left[(k_C D_2 + k_T D_1)\right] - \gamma N_2, \tag{4d}$$
$$\frac{dD_2}{dt} = D_0 H^{S-}(I_2, \lambda_{I_2,D_2}) - k_C D_2 N_2 - k_T N_1 D_2 - \gamma D_2, \tag{4e}$$
$$\frac{dI_2}{dt} = k_T N_2 D_1 - \gamma_I I_2, \tag{4f}$$

where $N_1$,$D_1$, and $I_1$ represent the Notch receptor, Delta, Jagged and NICD respectively for cell#1 and similarly $N_2$,$D_2$, and $I_2$ represent the Notch receptor, Delta, Jagged and NICD respectively for cell#2.

###### S1.2.2 Text S1.2.2 Notch-Delta-Jagged Module:

Similarly, the dynamics for two cells interacting with each other through NDJ signaling can be described as follows:

$$\frac{dN_1}{dt} = N_0 H^{S+}(I_1, \lambda_{I_1,N_1}) - N_1 \left[ (k_C D_1 + k_T D_2)\, H^{S+}(I_1, \lambda_{F,D}) + (k_C J_1 + k_T J_2)\, H^{S-}(I_1, \lambda_{F,J}) \right] - \gamma N_1, \tag{5a}$$

$$\frac{dD_1}{dt} = D_0 H^{S-}(I_1, \lambda_{I_1,D_1}) - k_C H^{S+}(I_1, \lambda_{F,D}) D_1 N_1 - k_T H^{S+}(I_2, \lambda_{F,D}) N_2 D_1 - \gamma D_1, \tag{5b}$$

$$\frac{dJ_1}{dt} = J_0 H^{S+}(I_1, \lambda_{I_1,J_1}) - k_C H^{S-}(I_1, \lambda_{F,J}) J_1 N_1 - k_T H^{S-}(I_2, \lambda_{F,J}) N_2 J_1 - \gamma J_1, \tag{5c}$$

$$\frac{dI_1}{dt} = k_T N_1 \left[ J_2 H^{S-}(I_1, \lambda_{F,J}) + D_2 H^{S+}(I_1, \lambda_{F,D}) \right] - \gamma_I I_1, \tag{5d}$$

$$\frac{dN_2}{dt} = N_0 H^{S+}(I_2, \lambda_{I_2,N_2}) - N_2 \left[ (k_C D_2 + k_T D_1)\, H^{S+}(I_2, \lambda_{F,D}) + (k_C J_2 + k_T J_1)\, H^{S-}(I_2, \lambda_{F,J}) \right] - \gamma N_2, \tag{5e}$$

$$\frac{dD_2}{dt} = D_0 H^{S-}(I_2, \lambda_{I_2,D_2}) - k_C H^{S+}(I_2, \lambda_{F,D}) D_2 N_2 - k_T H^{S+}(I_1, \lambda_{F,D}) N_1 D_2 - \gamma D_2, \tag{5f}$$

$$\frac{dJ_2}{dt} = J_0 H^{S+}(I_2, \lambda_{I_2,J_2}) - k_C H^{S-}(I_2, \lambda_{F,J}) J_2 N_2 - k_T H^{S-}(I_1, \lambda_{F,J}) N_1 J_2 - \gamma J_2, \tag{5g}$$

$$\frac{dI_2}{dt} = k_T N_2 \left[ J_1 H^{S-}(I_2, \lambda_{F,J}) + D_1 H^{S+}(I_2, \lambda_{F,D}) \right] - \gamma_I I_2, \tag{5h}$$

where $N_1$,$D_1$,$J_1$ and $I_1$ represent the Notch receptor, Delta, Jagged and NICD respectively for cell #1 and similarly $N_2$,$D_2$,$J_2$ and $I_2$ represent the Notch receptor, Delta, Jagged and NICD respectively for cell #2.

### S2 Text S2. Stochastic Models:

#### *S2.1* ***Text S2.1 One-Cell Model Equations:***

##### S2.1.1 Text S2.1.1 Notch-Delta Circuit:

We developed a stochastic model for eq (3), which corresponds the above deterministic model. In particular, we developed a definite form of master equation considering all the basic birth-death processes associated with the deterministic model and derived the Fokker-Planck equation from the master equation. We define $(N, D, I)^t$ as the state vector of the system. The transition probabilities along with all the molecular events those happen for this circuit are described in the following table.

**Table 1.** Eight different reactions for the Notch-Delta model (3), change of state vectors, gain and loss probabilities, and their propensity function. $V$ is the volume in which all the reactions occur. The symbols (+1) and (−1) in the column of state vectors represent birth and death processes of the respective chemical species. Here, $P$ stands for the grand probability function.

| Sl. No. | Elementary Events | Before Reaction | After Reaction | Gain Probability | Loss Probability | Propensity Function ($a_n$) |
|---|---|---|---|---|---|---|
| 1. | $\phi \xrightarrow{N_0 H^{S+}} N$ | $\begin{bmatrix} N-1 \\ D \\ I \end{bmatrix}$ | $\begin{bmatrix} N \\ D \\ I \end{bmatrix}$ | $VN_0P(N-1,D,I)$ | $VN_0P(N,D,I)$ | $VN_0$ |
| 2. | $\phi \xrightarrow{D_0 H^{S-}} D$ | $\begin{bmatrix} N \\ D-1 \\ I \end{bmatrix}$ | $\begin{bmatrix} N \\ D \\ I \end{bmatrix}$ | $VD_0P(N,D-1,I)$ | $VD_0P(N,D,I)$ | $VD_0$ |
| 3. | $N \xrightarrow{\gamma} \phi$ | $\begin{bmatrix} N+1 \\ D \\ I \end{bmatrix}$ | $\begin{bmatrix} N \\ D \\ I \end{bmatrix}$ | $\gamma(N+1)$ $\times P(N+1,D,I)$ | $\gamma NP(N,D,I)$ | $\gamma N$ |
| 4. | $D \xrightarrow{\gamma} \phi$ | $\begin{bmatrix} N \\ D+1 \\ I \end{bmatrix}$ | $\begin{bmatrix} N \\ D \\ I \end{bmatrix}$ | $\gamma(D+1)$ $\times P(N,D+1,I)$ | $\gamma DP(N,D,I)$ | $\gamma D$ |
| 5. | $I \xrightarrow{\gamma_I} \phi$ | $\begin{bmatrix} N \\ D \\ I+1 \end{bmatrix}$ | $\begin{bmatrix} N \\ D \\ I \end{bmatrix}$ | $\gamma_I(I+1)$ $\times P(N,D,I+1)$ | $\gamma_I IP(N,D,I)$ | $\gamma I$ |
| 6. | $D + N_{ext} \xrightarrow{k_T} \phi$ | $\begin{bmatrix} N \\ D+1 \\ I \end{bmatrix}$ | $\begin{bmatrix} N \\ D \\ I \end{bmatrix}$ | $k_T(D+1)N_{ext}$ $\times P(N,D+1,I)$ | $k_TDN_{ext}$ $\times P(N,D,I)$ | $k_TDN_{ext}$ |
| 7. | $N + D_{ext} \xrightarrow{k_T} I$ | $\begin{bmatrix} N+1 \\ D \\ I-1 \end{bmatrix}$ | $\begin{bmatrix} N \\ D \\ I \end{bmatrix}$ | $k_T(N+1)D_{ext}$ $\times P(N+1,D,I-1)$ | $k_TND_{ext}$ $\times P(N,D,I)$ | $k_TND_{ext}$ |
| 8. | $D + N \xrightarrow{k_C} \phi$ | $\begin{bmatrix} N+1 \\ D+1 \\ I \end{bmatrix}$ | $\begin{bmatrix} N \\ D \\ I \end{bmatrix}$ | $\frac{k_C}{V}(D+1)(N+1)$ $\times P(N+1,D+1,I)$ | $\frac{k_C}{V}DN$ $\times P(N,D,I)$ | $\frac{k_C}{V}DN$ |

Considering all the above fundamental processes, gain probability and loss probability, the corresponding master equation for the stochastic system is as follows:

$$
\begin{aligned}
\frac{\partial P(N,D,I)}{\partial t} &= VN_0P(N-1,D,I) - VN_0P(N,D,I) + VD_0P(N,D-1,I) - VD_0P(N,D,I) \\
&+ \gamma(N+1)P(N+1,D,I) - \gamma NP(N,D,I) + \gamma(D+1)P(N,D+1,I) \\
&- \gamma DP(N,D,I) + \gamma_I(I+1)P(N,D,I+1) - \gamma_I IP(N,D,I) \\
&+ k_T(D+1)(N_{ext})P(N,D+1,I) - k_TD(N_{ext})P(N,D,I) \\
&+ k_T(N+1)(D_{ext})P(N+1,D,I) - k_TN(D_{ext})P(N,D,I) \\
&+ \frac{k_C}{V}(D+1)(N+1)P(N+1,D+1,I) - \frac{k_C}{V}DNP(N,D,I)
\end{aligned}
$$

This master equation can be written using operator form, and this will be like:

$$
\begin{aligned}
\frac{\partial P(N,D,I,t)}{\partial t} &= [VN_0(E_N^{-1}-1) + VD_0(E_D^{-1}-1) + \gamma(E_N-1)N + \gamma(E_D-1)D + \gamma_I(E_I-1)I \\
&+ k_TN_{ext}(E_D-1)D + k_T(D_{ext})(E_NE_I^{-1}-1)N + \frac{k_C}{V}(E_DE_N-1)DN]P(N,D,I)
\end{aligned}
$$

Therefore the Fokker-Planck equation corresponding to the Notch-Delta-Jagged system is as follows:

$$
\begin{aligned}
\frac{\partial \rho}{\partial t} &= \frac{\partial}{\partial c_N}\left(-N_0 + \frac{k_T}{V}D_{ext}N + \frac{k_C}{V^2}DN + \frac{\gamma}{V}N\right)\rho + \frac{\partial}{\partial c_D}\left(-D_0 + \frac{k_C}{V^2}DN + \frac{k_T}{V}N_{ext}D + \frac{\gamma}{V}D\right)\rho \\
&+ \frac{\partial}{\partial c_I}\left(-\frac{k_T}{V}(D_{ext})N + \frac{\gamma_I}{V}I\right)\rho + \frac{\partial^2}{\partial c_N^2}\left(\frac{N_0}{2V} + \frac{k_T}{2V^2}(D_{ext})N + \frac{k_C}{2V^3}DN + \frac{\gamma}{2V^2}N\right)\rho \\
&+ \frac{\partial^2}{\partial c_D^2}(\frac{D_0}{2V} + \frac{k_C}{2V^3}DN + \frac{k_T}{2V^2}N_{ext}D + \frac{\gamma}{2V^2}D)\rho + \frac{\partial^2}{\partial c_I^2}\left(\frac{k_T}{2V^2}(D_{ext})N + \frac{\gamma_I}{2V^2}I\right)\rho \\
&- \frac{\partial}{\partial c_N}\frac{\partial}{\partial c_I}\left(\frac{k_T}{V^2}(D_{ext})N\right)\rho + \frac{\partial}{\partial c_N}\frac{\partial}{\partial c_D}\left(\frac{k_C}{V^3}DN\right)\rho
\end{aligned}
$$

##### S2.1.2 Text S2.1.2 Notch-Delta-Jagged Circuit:

**Table 2.** Twelve different reactions for the Notch-Delta-Jagged model (1), change of state vectors, gain and loss probabilities, and their propensity function. $V$ is the volume in which all the reactions occur. The symbols $(+1)$ and $(-1)$ in the column of state vectors represent birth and death processes of the respective chemical species. Here, $P$ stands for the grand probability function.

| Sl. No. | Elementary Events | Before Reaction | After Reaction | Gain Probability | Loss Probability | Propensity Function ($a_n$) |
|---|---|---|---|---|---|---|
| 1. | $\phi \xrightarrow{N_0 H^{S+}} N$ | $\begin{bmatrix} N-1 \\ D \\ J \\ I \end{bmatrix}$ | $\begin{bmatrix} N \\ D \\ J \\ I \end{bmatrix}$ | $VN_0P(N-1, D, J, I)$ | $VN_0P(N, D, J, I)$ | $VN_0$ |
| 2. | $\phi \xrightarrow{D_0 H^{S-}} D$ | $\begin{bmatrix} N \\ D-1 \\ J \\ I \end{bmatrix}$ | $\begin{bmatrix} N \\ D \\ J \\ I \end{bmatrix}$ | $VD_0P(N, D-1, J, I)$ | $VD_0P(N, D, J, I)$ | $VD_0$ |
| 3. | $\phi \xrightarrow{J_0 H^{S+}} J$ | $\begin{bmatrix} N \\ D \\ J-1 \\ I \end{bmatrix}$ | $\begin{bmatrix} N \\ D \\ J \\ I \end{bmatrix}$ | $VJ_0P(N, D, J-1, I)$ | $VJ_0P(N, D, J, I)$ | $VJ_0$ |
| 4. | $N \xrightarrow{\gamma} \phi$ | $\begin{bmatrix} N+1 \\ D \\ J \\ I \end{bmatrix}$ | $\begin{bmatrix} N \\ D \\ J \\ I \end{bmatrix}$ | $\gamma(N+1)$ $\times P(N+1, D, J, I)$ | $\gamma NP(N, D, J, I)$ | $\gamma N$ |
| 5. | $D \xrightarrow{\gamma} \phi$ | $\begin{bmatrix} N \\ D+1 \\ J \\ I \end{bmatrix}$ | $\begin{bmatrix} N \\ D \\ J \\ I \end{bmatrix}$ | $\gamma(D+1)$ $\times P(N, D+1, J, I)$ | $\gamma DP(N, D, J, I)$ | $\gamma D$ |
| 6. | $J \xrightarrow{\gamma} \phi$ | $\begin{bmatrix} N \\ D \\ J+1 \\ I \end{bmatrix}$ | $\begin{bmatrix} N \\ D \\ J \\ I \end{bmatrix}$ | $\gamma(J+1)$ $\times P(N, D, J+1, I)$ | $\gamma JP(N, D, J, I)$ | $\gamma J$ |

| Sl. No. | Elementary Events | Before Reaction | After Reaction | Gain Probability | Loss Probability | Propensity Function ($a_n$) |
|---|---|---|---|---|---|---|
| 7. | $N + (D_{ext} + J_{ext}) \xrightarrow{k_T} I$ | $\begin{bmatrix} N+1 \\ D \\ J \\ I-1 \end{bmatrix}$ | $\begin{bmatrix} N \\ D \\ J \\ I \end{bmatrix}$ | $k_T(N+1)(D_{ext}+J_{ext})$ $\times P(N+1, D, J, I-1)$ | $k_T N(D_{ext}+J_{ext})$ $\times P(N, D, J, I)$ | $k_T N$ $\times(D_{ext}+J_{ext})$ |
| 8. | $I \xrightarrow{\gamma_I} \phi$ | $\begin{bmatrix} N \\ D \\ J \\ I+1 \end{bmatrix}$ | $\begin{bmatrix} N \\ D \\ J \\ I \end{bmatrix}$ | $\gamma_I(I+1)$ $\times P(N, D, J, I+1)$ | $\gamma_I I P(N, D, J, I)$ | $\gamma_I I$ |
| 9. | $J + N \xrightarrow{k_C} \phi$ | $\begin{bmatrix} N+1 \\ D \\ J+1 \\ I \end{bmatrix}$ | $\begin{bmatrix} N \\ D \\ J \\ I \end{bmatrix}$ | $\frac{k_C}{V}(J+1)(N+1)$ $\times P(N+1, D, J+1, I)$ | $\frac{k_C}{V} JN$ $\times P(N, D, J, I)$ | $\frac{k_C}{V} JN$ |
| 10. | $D + N \xrightarrow{k_C} \phi$ | $\begin{bmatrix} N+1 \\ D+1 \\ J \\ I \end{bmatrix}$ | $\begin{bmatrix} N \\ D \\ J \\ I \end{bmatrix}$ | $\frac{k_C}{V}(D+1)(N+1)$ $\times P(N+1, D+1, J, I)$ | $\frac{k_C}{V} DN$ $\times P(N, D, J, I)$ | $\frac{k_C}{V} DN$ |
| 11. | $D + N_{ext} \xrightarrow{k_T} \phi$ | $\begin{bmatrix} N \\ D+1 \\ J \\ I \end{bmatrix}$ | $\begin{bmatrix} N \\ D \\ J \\ I \end{bmatrix}$ | $k_T(D+1)N_{ext}$ $\times P(N, D+1, J, I)$ | $k_T D N_{ext}$ $\times P(N, D, J, I)$ | $k_T D N_{ext}$ |
| 12. | $J + N_{ext} \xrightarrow{k_T} \phi$ | $\begin{bmatrix} N \\ D \\ J+1 \\ I \end{bmatrix}$ | $\begin{bmatrix} N \\ D \\ J \\ I \end{bmatrix}$ | $k_T(J+1)N_{ext}$ $\times P(N, D, J+1, I)$ | $k_T J N_{ext}$ $\times P(N, D, J, I)$ | $k_T J N_{ext}$ |

Considering all the above fundamental processes, gain probability and loss probability, the corresponding master equation for the stochastic system can be written as follows:

$$\frac{\partial P(N,D,J,I,t)}{\partial t} = VN_0P(N-1,D,J,I) - VN_0P(N,D,J,I) + VD_0P(N,D-1,J,I) - VD_0P(N,D,J,I)$$
$$+VJ_0P(N,D,J-1,I) - VJ_0P(N,D,J,I) + \gamma(N+1)P(N+1,D,J,I) - \gamma NP(N,D,J,I)$$
$$+\gamma(D+1)P(N,D+1,J,I) - \gamma DP(N,D,J,I) + \gamma(J+1)P(N,D,J+1,I) - \gamma JP(N,D,J,I)$$
$$+k_T(N+1)(D_{ext}+J_{ext})P(N+1,D,J,I-1) - k_TN(D_{ext}+J_{ext})P(N,D,J,I) + \gamma_I(I+1)P(N,D,J,I+1)$$
$$-\gamma_I IP(N,D,J,I) + \frac{k_C}{V}(J+1)(N+1)P(N+1,D,J+1,I) - \frac{k_C}{V}JNP(N,D,J,I)$$
$$+\frac{k_C}{V}(D+1)(N+1)P(N+1,D+1,J,I) - \frac{k_C}{V}DNP(N,D,J,I) + k_T(D+1)N_{ext}P(N,D+1,J,I)$$
$$-k_TDN_{ext}P(N,D,J,I) + k_T(J+1)N_{ext}P(N,D,J+1,I) - k_TJN_{ext}P(N,D,J,I)$$

The above equation can be written in the form of a shift operator, then the equation becomes:

$$\begin{aligned}\frac{\partial P(N,D,J,I,t)}{\partial t} &= [VN_0(E_N^{-1}-1) + VD_0(E_D^{-1}-1) + VJ_0(E_J^{-1}-1) + \gamma(E_N-1)N + \gamma(E_D-1)D \\ &+ \gamma(E_J-1)J + k_T(D_{ext}+J_{ext})(E_NE_I^{-1}-1)N + \gamma_I(E_I-1)I \\ &+ \frac{k_C}{V}(E_JE_N-1)JN + \frac{k_C}{V}(E_DE_N-1)DN \\ &+ k_TN_{ext}(E_D-1)D + k_TN_{ext}(E_J-1)J]P(N,D,J,I)\end{aligned}$$

Thus, the Fokker-Planck equation can be obtained as:

$$
\begin{aligned}
\frac{\partial \rho}{\partial t} = & \frac{\partial}{\partial c_N}\left(-N_0 + \frac{k_T}{V}(D_{ext}+J_{ext})N + \frac{k_C}{V^2}JN + \frac{k_C}{V^2}DN + \frac{\gamma}{V}N\right)\rho \\
& + \frac{\partial}{\partial c_D}\left(-D_0 + \frac{k_C}{V^2}DN + \frac{k_T}{V}N_{ext}D + \frac{\gamma}{V}D\right)\rho + \frac{\partial}{\partial c_J}\left(-J_0 + \frac{k_C}{V^2}JN + \frac{k_T}{V}N_{ext}J + \frac{\gamma}{V}J\right)\rho \\
& + \frac{\partial}{\partial c_I}\left(-\frac{k_T}{V}(D_{ext}+J_{ext})N + \frac{\gamma_I}{V}I\right)\rho \\
& + \frac{\partial^2}{\partial c_N^2}\left(\frac{N_0}{2V} + \frac{k_T}{2V^2}(D_{ext}+J_{ext})N + \frac{k_C}{2V^3}JN + \frac{k_C}{2V^3}DN + \frac{\gamma}{2V^2}N\right)\rho \\
& + \frac{\partial^2}{\partial c_D^2}(\frac{D_0}{2V} + \frac{k_C}{2V^3}DN + \frac{k_T}{2V^2}N_{ext}D \\
& + \frac{\gamma}{2V^2}D)\rho + \frac{\partial^2}{\partial c_J^2}\left(\frac{J_0}{2V} + \frac{k_C}{2V^3}JN + \frac{k_T}{2V^2}N_{ext}J + \frac{\gamma}{2V^2}J\right)\rho \\
& + \frac{\partial^2}{\partial c_I^2}\left(\frac{k_T}{2V^2}(D_{ext}+J_{ext})N + \frac{\gamma_I}{2V^2}I\right)\rho \\
& - \frac{\partial}{\partial c_N}\frac{\partial}{\partial c_I}\left(\frac{k_T}{V^2}(D_{ext}+J_{ext})N\right)\rho + \frac{\partial}{\partial c_N}\frac{\partial}{\partial c_J}\left(\frac{k_C}{V^3}JN\right)\rho + \frac{\partial}{\partial c_N}\frac{\partial}{\partial c_D}\left(\frac{k_C}{V^3}DN\right)\rho
\end{aligned}
$$

##### S2.1.3 Text S2.1.3 Fringe mediated Notch-Delta-Jagged Circuit:

We developed a stochastic formulation corresponding to the deterministic model given in eq (2). Specifically, we constructed an explicit master equation by accounting for all fundamental birth–death processes associated with the deterministic dynamics, and subsequently derived the corresponding Fokker–Planck equation from the master equation. The state of the system is represented by the vector $(N, D, I)^{\mathsf{T}}$. The transition probabilities, together with the underlying molecular events governing the dynamics of this circuit, are summarized in the table below. The elementary microscopic processes constituting the Fringe-mediated Notch–Delta–Jagged circuit described by Eq. (2) are listed as follows:

**Table 3.** Twelve different reactions for the Notch-Delta-Jagged model ((**??**)), change of state vectors, gain and loss probabilities, and their propensity function. $V$ is the volume in which all the reactions occur. The symbols $(+1)$ and $(-1)$ in the column of state vectors represent birth and death processes of the respective chemical species. Here, $P$ stands for the grand probability function.

| Sl. No. | Elementary Events | Before Reaction | After Reaction | Gain Probability | Loss Probability | Propensity Function ($a_n$) |
|---|---|---|---|---|---|---|
| 1. | $\phi \xrightarrow{N_0 H^{S+}} N$ | $\begin{bmatrix} N-1 \\ D \\ J \\ I \end{bmatrix}$ | $\begin{bmatrix} N \\ D \\ J \\ I \end{bmatrix}$ | $VN_0P(N-1, D, J, I)$ | $VN_0P(N, D, J, I)$ | $VN_0$ |
| 2. | $\phi \xrightarrow{D_0 H^{S-}} D$ | $\begin{bmatrix} N \\ D-1 \\ J \\ I \end{bmatrix}$ | $\begin{bmatrix} N \\ D \\ J \\ I \end{bmatrix}$ | $VD_0P(N, D-1, J, I)$ | $VD_0P(N, D, J, I)$ | $VD_0$ |
| 3. | $\phi \xrightarrow{J_0 H^{S+}} J$ | $\begin{bmatrix} N \\ D \\ J-1 \\ I \end{bmatrix}$ | $\begin{bmatrix} N \\ D \\ J \\ I \end{bmatrix}$ | $VJ_0P(N, D, J-1, I)$ | $VJ_0P(N, D, J, I)$ | $VJ_0$ |
| 4. | $N \xrightarrow{\gamma} \phi$ | $\begin{bmatrix} N+1 \\ D \\ J \\ I \end{bmatrix}$ | $\begin{bmatrix} N \\ D \\ J \\ I \end{bmatrix}$ | $\gamma(N+1)$ $\times P(N+1, D, J, I)$ | $\gamma NP(N, D, J, I)$ | $\gamma N$ |
| 5. | $D \xrightarrow{\gamma} \phi$ | $\begin{bmatrix} N \\ D+1 \\ J \\ I \end{bmatrix}$ | $\begin{bmatrix} N \\ D \\ J \\ I \end{bmatrix}$ | $\gamma(D+1)$ $\times P(N, D+1, J, I)$ | $\gamma DP(N, D, J, I)$ | $\gamma D$ |
| 6. | $J \xrightarrow{\gamma} \phi$ | $\begin{bmatrix} N \\ D \\ J+1 \\ I \end{bmatrix}$ | $\begin{bmatrix} N \\ D \\ J \\ I \end{bmatrix}$ | $\gamma(J+1)$ $\times P(N, D, J+1, I)$ | $\gamma JP(N, D, J, I)$ | $\gamma J$ |

| Sl. No. | Elementary Events | Before Reaction | After Reaction | Gain Probability | Loss Probability | Propensity Function ($a_n$) |
|---|---|---|---|---|---|---|
| 7. | $N + D_{ext} \xrightarrow{k_{T_D}} I$ | $\begin{bmatrix} N+1 \\ D \\ J \\ I-1 \end{bmatrix}$ | $\begin{bmatrix} N \\ D \\ J \\ I \end{bmatrix}$ | $k_{T_D}(N+1)D_{ext} \times P(N+1, D, J, I-1)$ | $k_{T_D} N D_{ext} \times P(N, D, J, I)$ | $k_{T_D} N \times D_{ext}$ |
| 8. | $I \xrightarrow{\gamma_I} \phi$ | $\begin{bmatrix} N \\ D \\ J \\ I+1 \end{bmatrix}$ | $\begin{bmatrix} N \\ D \\ J \\ I \end{bmatrix}$ | $\gamma_I(I+1) \times P(N, D, J, I+1)$ | $\gamma_I I P(N, D, J, I)$ | $\gamma_I I$ |
| 9. | $J + N \xrightarrow{k_{C_J}} \phi$ | $\begin{bmatrix} N+1 \\ D \\ J+1 \\ I \end{bmatrix}$ | $\begin{bmatrix} N \\ D \\ J \\ I \end{bmatrix}$ | $\frac{k_{C_J}}{V}(J+1)(N+1) \times P(N+1, D, J+1, I)$ | $\frac{k_{C_J}}{V} J N \times P(N, D, J, I)$ | $\frac{k_{C_J}}{V} J N$ |
| 10. | $D + N \xrightarrow{k_{C_D}} \phi$ | $\begin{bmatrix} N+1 \\ D+1 \\ J \\ I \end{bmatrix}$ | $\begin{bmatrix} N \\ D \\ J \\ I \end{bmatrix}$ | $\frac{k_{C_D}}{V}(D+1)(N+1) \times P(N+1, D+1, J, I)$ | $\frac{k_{C_D}}{V} D N \times P(N, D, J, I)$ | $\frac{k_{C_D}}{V} D N$ |
| 11. | $D + N_{ext} \xrightarrow{k_{T_D}} \phi$ | $\begin{bmatrix} N \\ D+1 \\ J \\ I \end{bmatrix}$ | $\begin{bmatrix} N \\ D \\ J \\ I \end{bmatrix}$ | $k_{T_D}(D+1)N_{ext} \times P(N, D+1, J, I)$ | $k_{T_D} D N_{ext} \times P(N, D, J, I)$ | $k_{T_D} D N_{ext}$ |
| 12. | $J + N_{ext} \xrightarrow{k_{T_J}} \phi$ | $\begin{bmatrix} N \\ D \\ J+1 \\ I \end{bmatrix}$ | $\begin{bmatrix} N \\ D \\ J \\ I \end{bmatrix}$ | $k_{T_J}(J+1)N_{ext} \times P(N, D, J+1, I)$ | $k_{T_J} J N_{ext} \times P(N, D, J, I)$ | $k_{T_J} J N_{ext}$ |
| 13. | $N + J_{ext} \xrightarrow{k_{T_J}} I$ | $\begin{bmatrix} N+1 \\ D \\ J \\ I-1 \end{bmatrix}$ | $\begin{bmatrix} N \\ D \\ J \\ I \end{bmatrix}$ | $k_{T_J}(N+1)J_{ext} \times P(N+1, D, J, I-1)$ | $k_{T_J} N J_{ext} \times P(N, D, J, I)$ | $k_{T_J} N \times J_{ext}$ |

#### *S2.2* ***Text S2.2 Two-Cell Model Equations:***

##### S2.2.1 Text S2.2.1 Notch-Delta Circuit:

For the deterministic model (4) describing two-cell Notch–Delta communication, the elementary reactions are given as follows:

**Table 4.** Different fundamental processes for the two-cell Notch-Delta model, their propensity function, and the corresponding jump of species. $V$ is the volume in which all the reactions occur. The symbols (+1) and (−1) in the column of jump of molecules represent birth and death processes of the respective chemical species.

| Sl. No. | Elementary Events | Propensity Function ($a_n$) | Jump of molecules |
|---|---|---|---|
| 1. | $\phi \xrightarrow{N_0 H^{S+}} N_i$ | $V N_0 H^{S+}$ | $N_i = N_i + 1$ |
| 2. | $\phi \xrightarrow{D_0 H^{S-}} D_i$ | $V D_0 H^{S-}$ | $D_i = D_i + 1$ |
| 3. | $N_i \xrightarrow{\gamma} \phi$ | $\gamma N_i$ | $N_i = N_i - 1$ |
| 4. | $D_i \xrightarrow{\gamma} \phi$ | $\gamma D_i$ | $D_i = D_i - 1$ |
| 5. | $I_i \xrightarrow{\gamma_I} \phi$ | $\gamma_{I_i} I_i$ | $I_i = I_i - 1$ |
| 6. | $D_i + N_i \xrightarrow{k_C} \phi$ | $k_C D_i N_i$ | $N_i = N_i - 1$<br>$D_i = D_i - 1$ |
| 7. | $N_i + D_j \xrightarrow{k_T} I_i, i \neq j$ | $k_T N_i D_j$ | $N_i = N_i - 1$<br>$D_j = D_j - 1$<br>$I_i = I_i + 1$ |

##### S2.2.2 Text S2.2.2 Notch-Delta-Jagged Circuit:

Similarly, the elementary equations for two cells interacting with each other through NDJ signaling (5) can be described as follows:

**Table 5.** Different fundamental processes for the two-cell Notch-Delta model, their propensity function, and the corresponding jump of species. $V$ is the volume in which all the reactions occur. The symbols $(+1)$ and $(-1)$ in the column of jump of molecules represent birth and death processes of the respective chemical species.

| Sl. No. | Elementary Events | Propensity Function ($a_n$) | Jump of molecules |
|---|---|---|---|
| 1. | $\phi \xrightarrow{N_0 H^{S+}} N_i$ | $V N_0 H^{S+}$ | $N_i = N_i + 1$ |
| 2. | $\phi \xrightarrow{D_0 H^{S-}} D_i$ | $V D_0 H^{S-}$ | $D_i = D_i + 1$ |
| 3. | $\phi \xrightarrow{J_0 H^{S+}} J_i$ | $V J_0 H^{S+}$ | $J_i = J_i + 1$ |
| 4. | $N_i \xrightarrow{\gamma} \phi$ | $\gamma N_i$ | $N_i = N_i - 1$ |
| 5. | $D_i \xrightarrow{\gamma} \phi$ | $\gamma D_i$ | $D_i = D_i - 1$ |
| 6. | $J_i \xrightarrow{\gamma} \phi$ | $\gamma J_i$ | $J_i = J_i - 1$ |
| 7. | $I_i \xrightarrow{\gamma_I} \phi$ | $\gamma I_i$ | $I_i = I_i - 1$ |
| 8. | $D_i + N_i \xrightarrow{k_C} \phi$ | $k_T D_i N_i$ | $N_i = N_i - 1$<br>$D_i = D_i - 1$ |
| 9. | $N_i + D_j \xrightarrow{k_T} I_i, i \neq j$ | $k_T N_i D_j$ | $N_i = N_i - 1$<br>$D_j = D_j - 1$<br>$I_i = I_i + 1$ |
| 10. | $J_i + N_i \xrightarrow{k_C} \phi$ | $k_T J_i N_i$ | $N_i = N_i - 1$<br>$j_i = j_i - 1$ |
| 11. | $N_i + J_j \xrightarrow{k_T} I_i, i \neq j$ | $k_T N_i J_j$ | $N_i = N_i - 1$<br>$J_j = J_j - 1$<br>$I_i = I_i + 1$ |

### S3 Text S3. Methodology:

#### *S3.1* ***Text S3.1 Numerical bifurcation diagrams and Monte Carlo simulations:***

All numerical bifurcation diagrams were obtained using the continuation package MATCONT. Deterministic simulations were performed in MATLAB (2018b) using the equations and parameter values reported in Boareto et al (2015). The original model files used to reproduce the bifurcation diagrams shown in Figures 2, 5, and 7, as well as Figures S1, S2, S3, S5, S7, S10, S12, and S13, are publicly available at (`https://github.com/mboareto/BoaretoEtAl2015PNAS`)

Stochastic trajectories of Notch levels were generated by simulating the corresponding master equation using Monte Carlo methods. The simulations in Figures 2,3,5, 7, 8, and in all supplementary figures were carried out using the Gillespie algorithm Gillespie (1977). The reaction events and all system parameters are presented in **SI**. Here, both the time and bifurcation parameter are varied together to obtain the time series. More details of the parameters for simulation are presented in the text of **SI**.

#### *S3.2* ***Text S3.2 Early Warning Indicators:***

In a multistable system, demographic noise can drive tipping events, which can be anticipated using critical slowing down (CSD)-based indicators. We calculated variance, autocorrelation, return rate, density ratio, skewness, and kurtosis to forecast impending transitions (Figures 2-5 and 7). Time series analyses were performed using the Early Warning Signals Toolbox (`http://www.earlywarning-signals.org/`). Further details of the methodology are presented in Sarkar et al (2019).

#### *S3.3* ***Text S3.3 Stochastic Potential Analysis:***

We calculated the stochastic potential (Figure 6 and Figures S4, S6, S8, S9, and S11) numerically using trajectories obtained from the stochastic simulations, following the method described in Sarkar et al (2019).

**Table 6. Parameter values used in the simulations**

ND, NDJ, and NDJF correspond to the systems described by Eq. (1), Eq. (2), and the Fringe-mediated version of Eq. (2), respectively mentioned in the main manuscript.

| Sl.no | parameter | Value | Value | Value |
|---|---|---|---|---|
| - | – | $ND$ | $NDJ$ | $NDJF$ |
| 1 | $\gamma$ | 0.1 | 0.1 | 0.1 |
| 2 | $\gamma_I$ | 0.5 | 0.5 | 0.5 |
| 3 | $N_0$ | 500 | 1600 | 1400 |
| 4 | $D_0$ | 1000 | 1800 | 1600 |
| 5 | $J_0$ | – | 1200 | 1200 |
| 6 | $k_T$ | 0.00005 | 0.00005 | 0.00005 or 0.000025 |
| 7 | $k_C$ | 0.0005 | 0.0005 | 0.0005 |
| 8 | $I_0$ | 200 | 200 | 200 |
| 9 | $n_N$ | 2.0 | 2.0 | 2.0 |
| 10 | $n_D$ | 2.0 | 2.0 | 2.0 |
| 11 | $n_J$ | – | 5.0 | 5.0 |
| 12 | $n_F$ | – | 1.0 | 1.0 |
| 13 | $\lambda_N$ | 2.0 | 2.0 | 2.0 |
| 14 | $\lambda_J$ | – | 2.0 | 2.0 |
| 15 | $\lambda_D$ | 0.0 | 0.0 | 0.0 |
| 16 | $\lambda_N^F$ | – | – | 3.0 or 1 |
| 17 | $\lambda_J^F$ | – | – | 0.3 or 1 |

### S4 Supplementary Figures

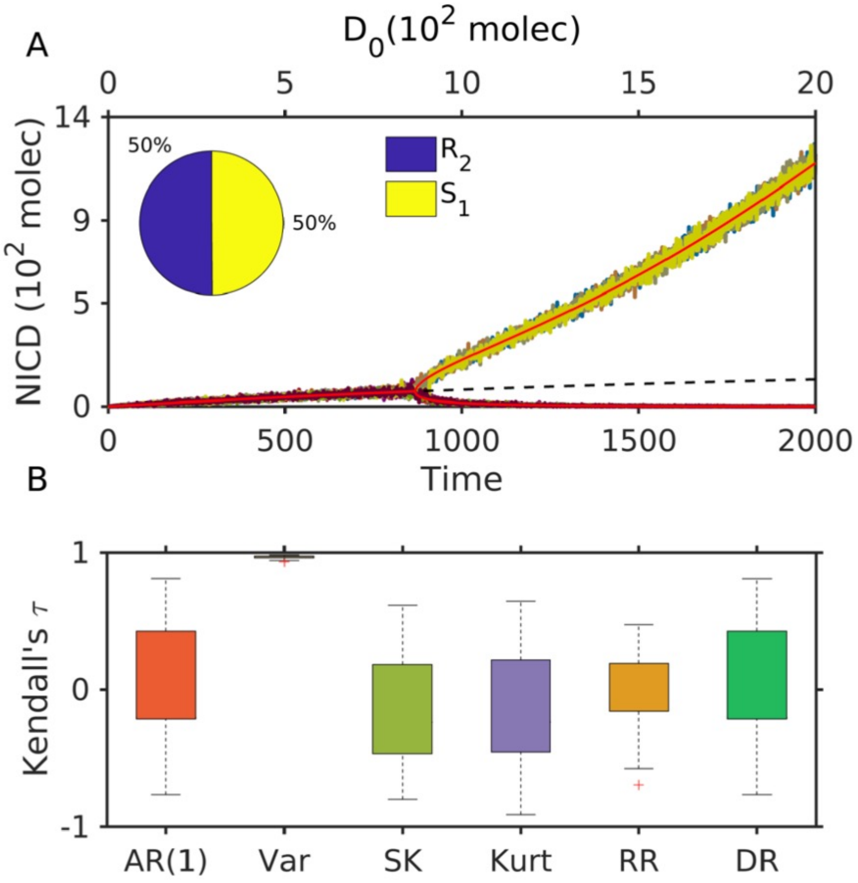

**Figure S1.** A) Bifurcation diagram and transitions between different cell states for two interacting ND cells. Solid (red) lines indicate stable steady states, and the dotted (black) line indicates the unstable steady state of the corresponding deterministic model. 100 replicates of stochastic time series are indicated by fluctuating lines. B) Performance of CSD indicators of 100 replicates of trajectories. Kendall-$\tau$ rank correlations obtained from replicates of trajectories of different CSD indicators, autocorrelation at lag-1 (AR(1)), variance (Var), skewness (SK), Kurtosis (Kurt), return rate (RR), and density ratio (DR). The box plots display the median, with the bottom and top edges of the box representing the 25th and 75th percentiles, respectively. The pie diagram in A shows a basin stability measure at $D_0 = 1000$. Box whiskers indicate the minimum and maximum values.

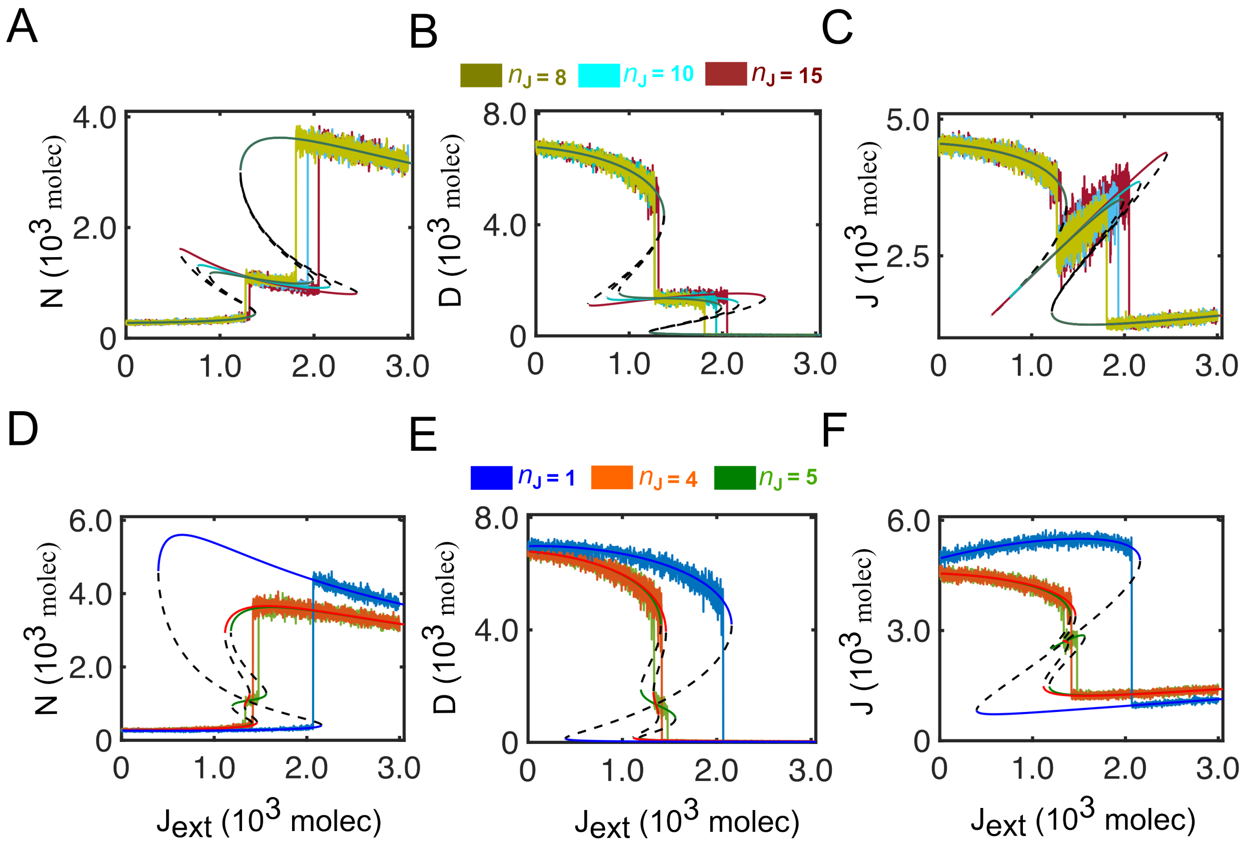

**Figure S2.** Bifurcation diagram and flickering of Notch (N), Delta (D), and Jagged (J) by varying $n_J$ for NDJ system without the effect of Fringe (system-1).

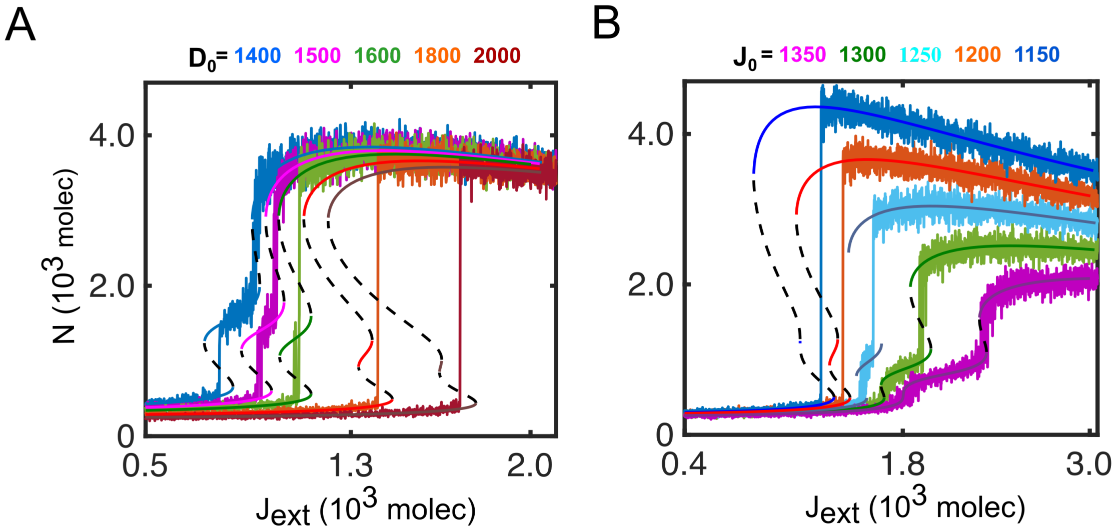

**Figure S3.** Bifurcation diagram and flickering of Notch (N) by varying A) the production rate of Delta $D_0$ and B) the production rate of Jagged $J_0$ for NDJ system (system-1) in case of $n_J = 4$.

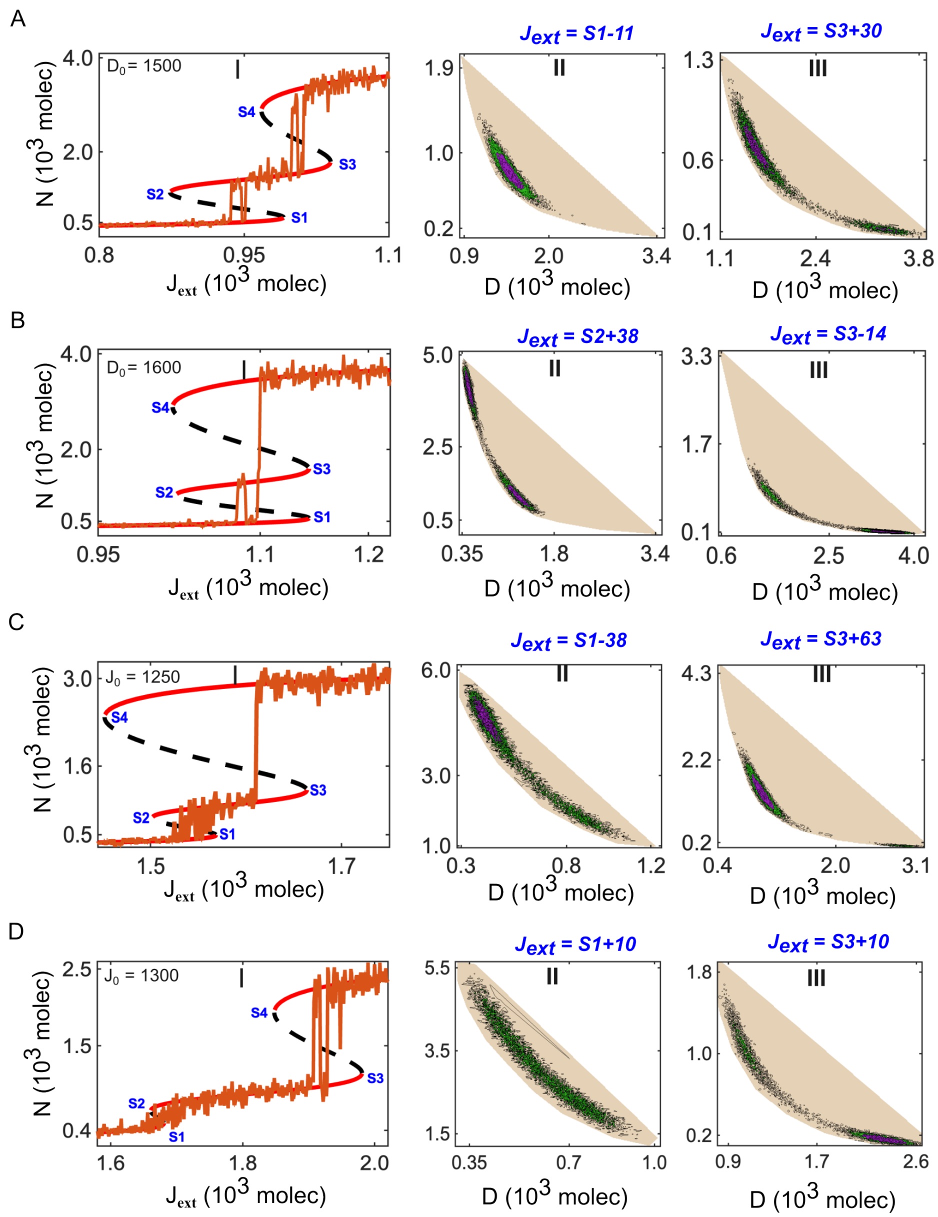

**Figure S4.** Potential landscape of the NDJ circuit (eq 1) under varying $J_{ext}$. (A) At Delta production rate of $D_0 = 1500$. (I) Bifurcation diagram showing the steady-state Notch level as a function of $J_{ext}$, exhibiting multiple saddle points (S1 = 991, S2 = 873, S3 = 1040, S4 = 968). Solid red curves denote stable equilibria, while dashed black curves denote unstable equilibria. (II, III) Stochastic potential landscapes for representative values of $J_{ext}$ corresponding to S1 (II) and S3 (III). Deeper colors indicate deeper potential wells (local minima) representing stable attractors. (B) At the Delta production rate of $D_0 = 1600$. (I) Bifurcation diagram of the steady-state Notch level versus $J_{ext}$, showing a qualitatively altered bifurcation structure (S1 = 1440, S2 = 1022, S3 = 1144, S4 = 1018). (II, III) Corresponding stochastic potential landscapes for different values of $J_{ext}$.(C) At Jagged production rate of $J_0 = 1250$. (I) Bifurcation diagram of the steady-state Notch level versus $J_{ext}$ (S1 = 1568, S2 =1501, S3=1663, S4=1451). (II, III) Stochastic potential landscapes for different values of $J_{ext}$. (D) At Jagged production rate of $J_0 = 1300$. (I) Bifurcation diagram of the steady-state Notch level versus $J_{ext}$, showing a qualitatively altered bifurcation structure (S1 = 1680, S2 =1660, S3=1980, S4=1847). (II, III) Under a higher Jagged production rate, flickering mediated uniform distribution of stochastic potential landscapes for different values of $J_{ext}$, illustrating the reshaping of basins attractor.

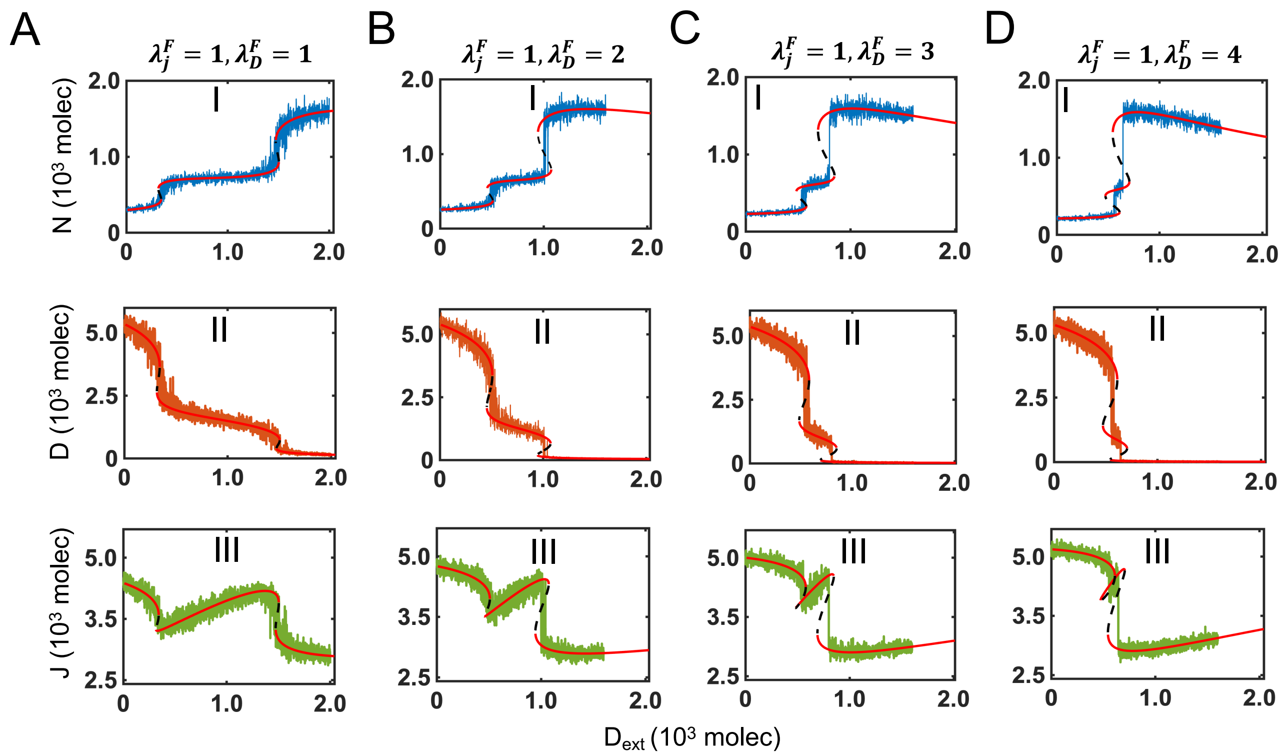

**Figure S5.** Bifurcation diagram fo steady state level of Notch, Delta and Jagged versus $D_{ext}$ and corresponding flickering for NDJ system ($nj = 5, WT$) under Fringe mediated effect on only Delta binding at (A) $\lambda_D^F = 1$ and $\lambda_J^F = 1$, (B) $\lambda_D^F = 2$ and $\lambda_J^F = 1$, (C) $\lambda_D^F = 3$ and $\lambda_J^F = 1$, and (D) $\lambda_D^F = 4$ and $\lambda_J^F = 1$.

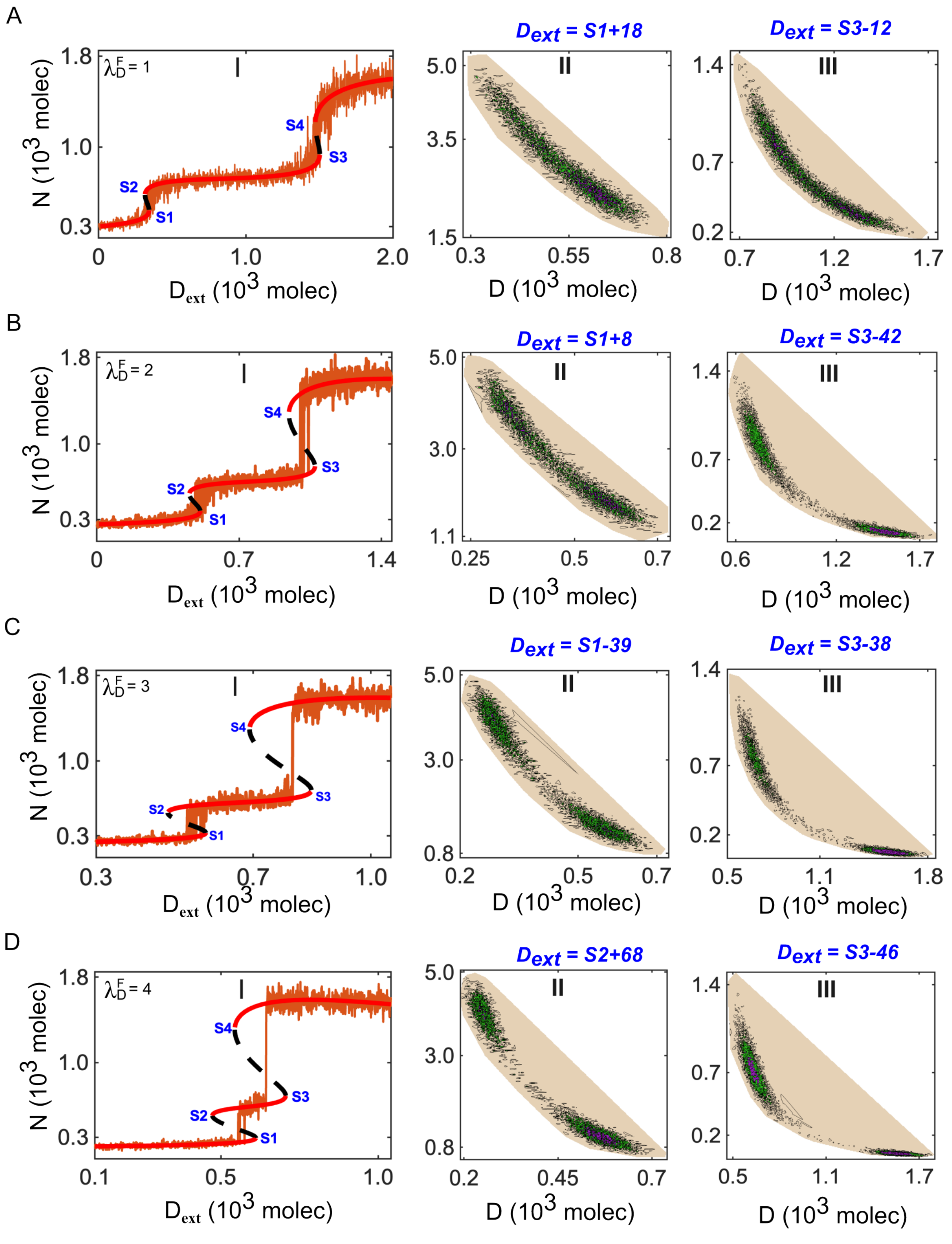

**Figure S6.** Potential landscape of the NDJ circuit system under Fringe mediated effect on only Notch-Delta binding ($n_j = 5$, WT) ($\lambda_D^F > 1$ and $\lambda_J^F = 1$) (A) Bifurcation diagram exhibiting multiple saddle points ($S1 = 342, S2 = 315, S3 = 1502, S4 = 1472$) and stochastic potential landscapes for representative values of $D_{ext}$ at $\lambda_D^F = 1$. (B) Bifurcation diagram exhibiting multiple saddle points ($S1 = 508, S2 = 454, S3 = 1072, S4 = 944$) and stochastic potential landscapes for representative values of $D_{ext}$ at $\lambda_D^F = 2$. (C) Bifurcation diagram exhibiting multiple saddle points ($S1 = 579, S2 = 482, S3 = 848, S4 = 690$) and stochastic potential landscapes for representative values of $D_{ext}$ at $\lambda_D^F = 3$. (D) Bifurcation diagram exhibiting multiple saddle points ($S1 = 611, S2 = 472, S3 = 706, S4 = 543$) and stochastic potential landscapes for representative values of $D_{ext}$ at $\lambda_D^F = 4$.

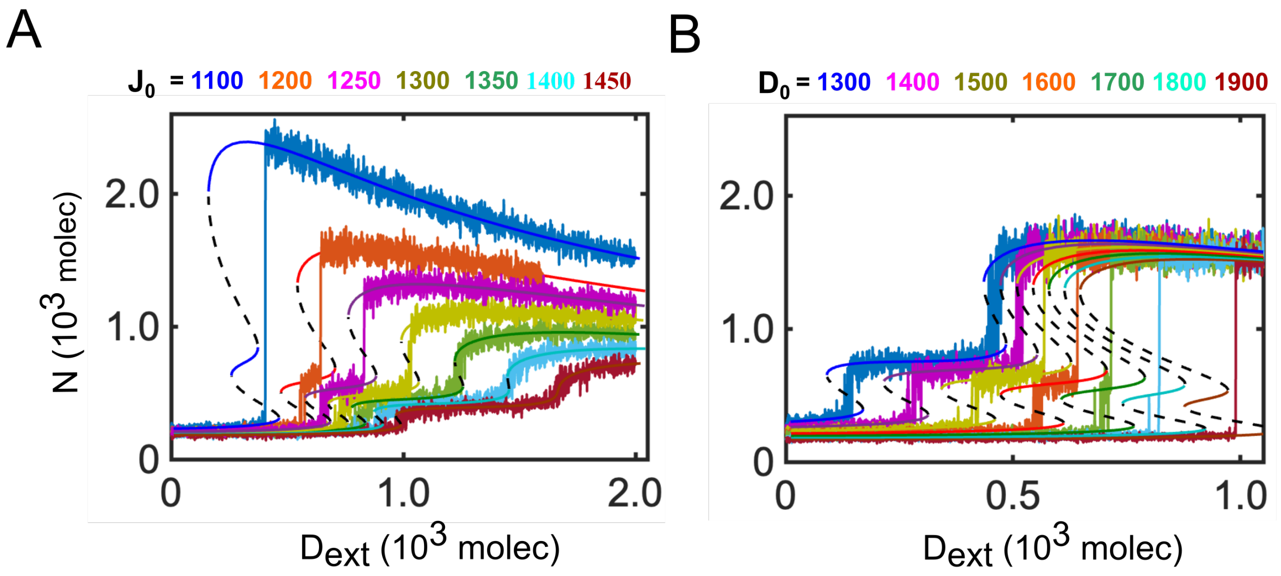

**Figure S7.** Bifurcation diagram and flickering of Notch (N) by varying A) the production rate of Jagged $J_0$ and B) the production rate of Delta $D_0$ for Fringe mediated NDJ system $\lambda_D^F = 4$, $\lambda_D^F = 1$.

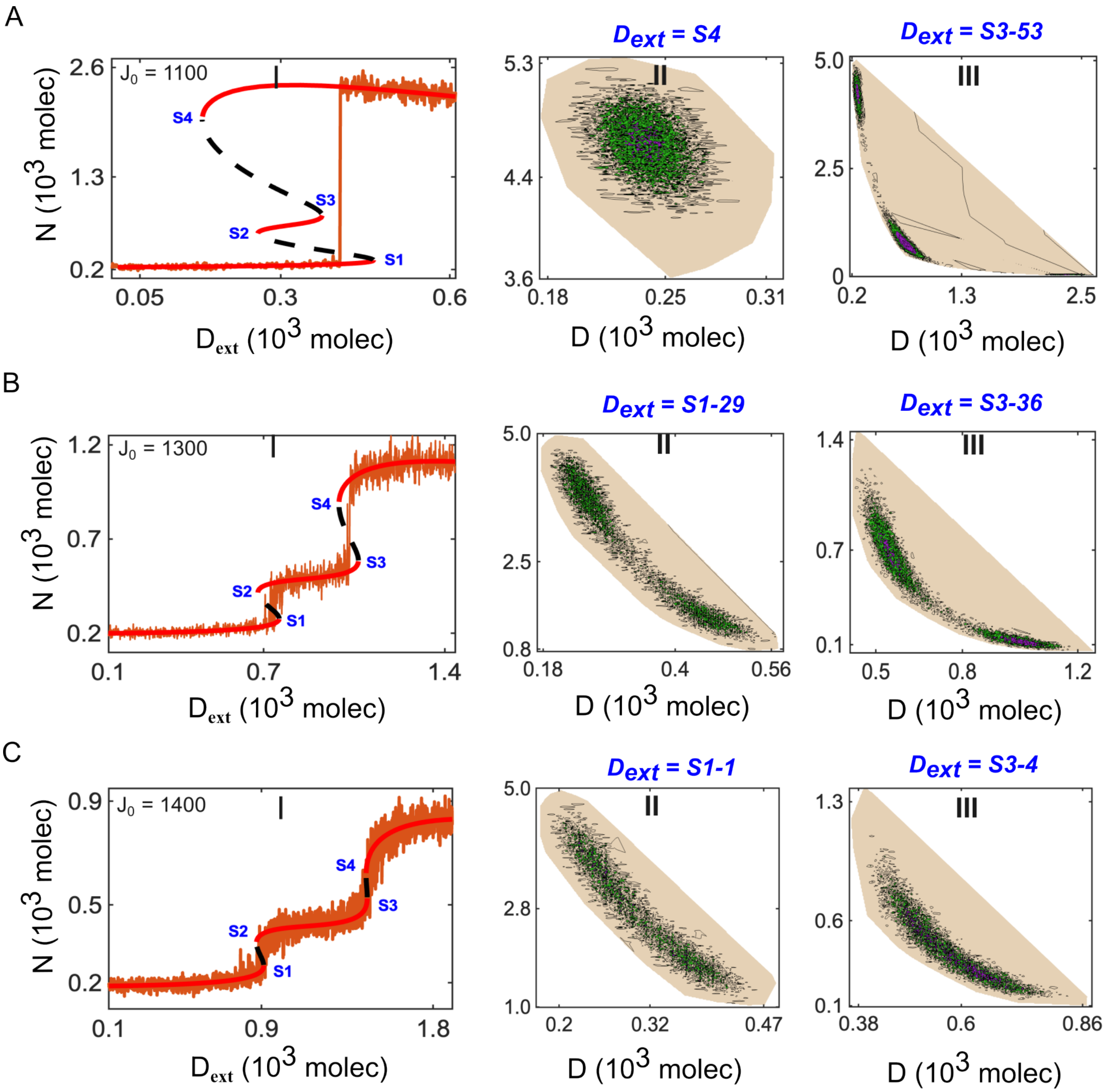

**Figure S8.** Potential landscape of the NDJ circuit system under strong Fringe mediated effect on Notch-Delta binding ($n_j = 5$, WT) ($\lambda_D^F = 4$ and $\lambda_J^F = 1$) at different levels of Jagged ($J_0$). (A) At Jagged production rate of $J_0 = 1100$. (I) Bifurcation diagram showing the steady-state Notch level as a function of $D_{ext}$, exhibiting multiple saddle points ($S1 = 464, S2 = 259, S3 = 373, S4 = 160$). (II, III) Stochastic potential landscapes for representative values of $D_{ext}$ corresponding to S4 (II) and S3 (III). Deeper colors indicate deeper potential wells (local minima) representing stable attractors. (B) At Jagged production rate of $J_0 = 1300$. (I) Bifurcation diagram showing the steady-state Notch level as a function of $D_{ext}$, exhibiting multiple saddle points ($S1 = 759, S2 = 678, S3 = 1066, S4 = 991$). (II, III) Stochastic potential landscapes for representative values of $D_{ext}$. (C) At Jagged production rate of $J_0 = 1400$. (I) Bifurcation diagram showing the steady-state Notch level as a function of $D_{ext}$, exhibiting multiple saddle points ($S1 = 911, S2 = 873, S3 = 1454, S4 = 1448$). Under higher Jagged production rate ($J_0 = 1400$), flickering mediated uniform distribution of stochastic potential for different values of $D_{ext}$, illustrating the reshaping of basins attractor.

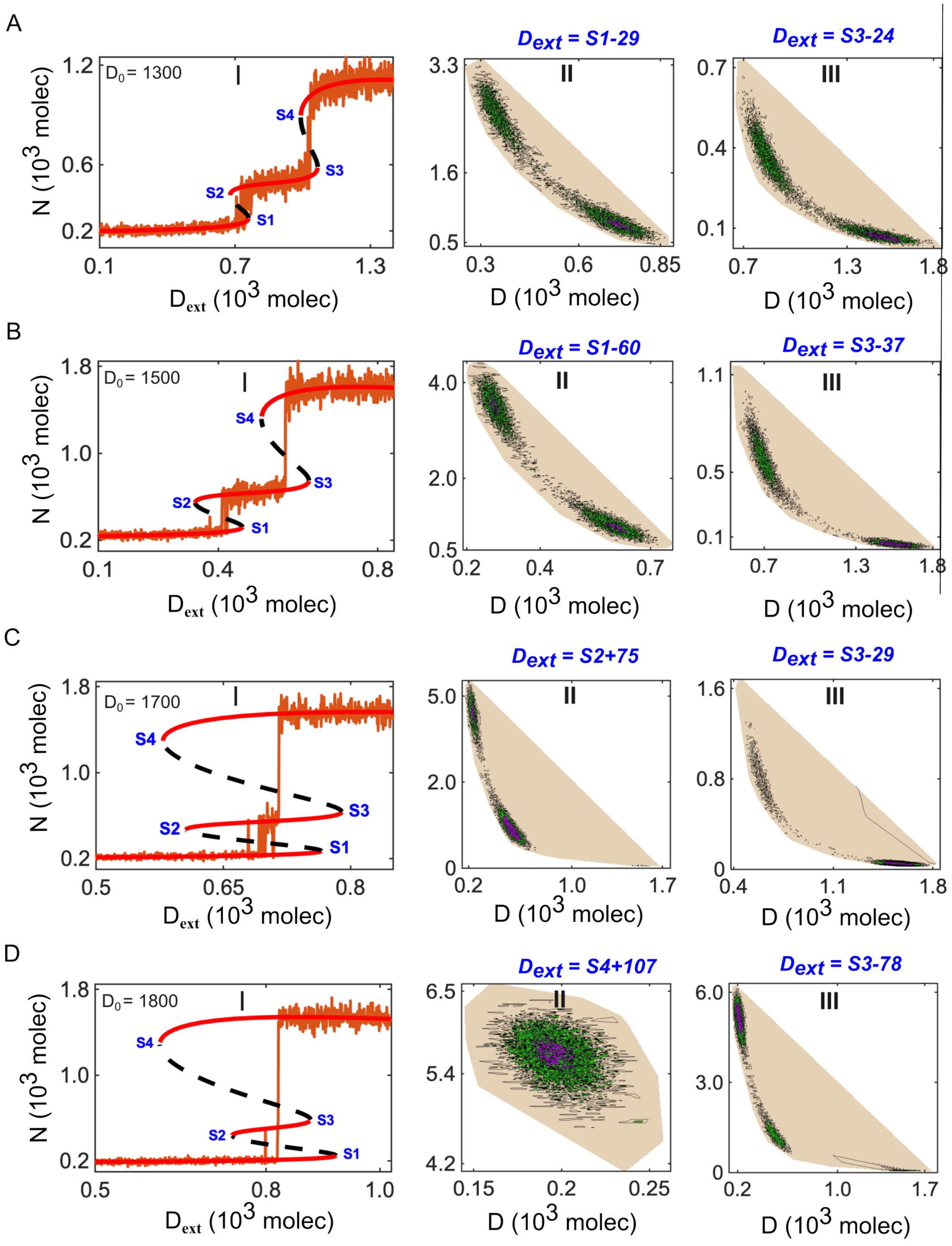

**Figure S9.** Potential landscape of the NDJ circuit system under strong Fringe mediated effect on Notch-Delta binding ($n_j$ = 5, WT) ($\lambda_D^F$ = 4 and $\lambda_J^F$ = 1) at different levels of Delta ($J_0$). (A) Bifurcation diagram exhibiting multiple saddle points ($S1 = 169, S2 = 90, S3 = 484, S4 = 435$) and stochastic potential landscapes for representative values of $D_{ext}$ at $D_0 = 1300$. (B) Bifurcation diagram exhibiting multiple saddle points ($S1 = 460, S2 = 342, S3 = 627, S4 = 508$) and stochastic potential landscapes for representative values of $D_{ext}$ at $D_0 = 1500$. (C) Bifurcation diagram exhibiting multiple saddle points ($S1 = 765, S2 = 605, S3 = 789, S4 = 578$) and stochastic potential landscapes for representative values of $D_{ext}$ at $D_0 = 1700$. (D) Bifurcation diagram exhibiting multiple saddle points ($S1 = 922, S2 = 740, S3 = 878, S4 = 543$) and stochastic potential landscapes for representative values of $D_{ext}$ at $D_0 = 1700$.

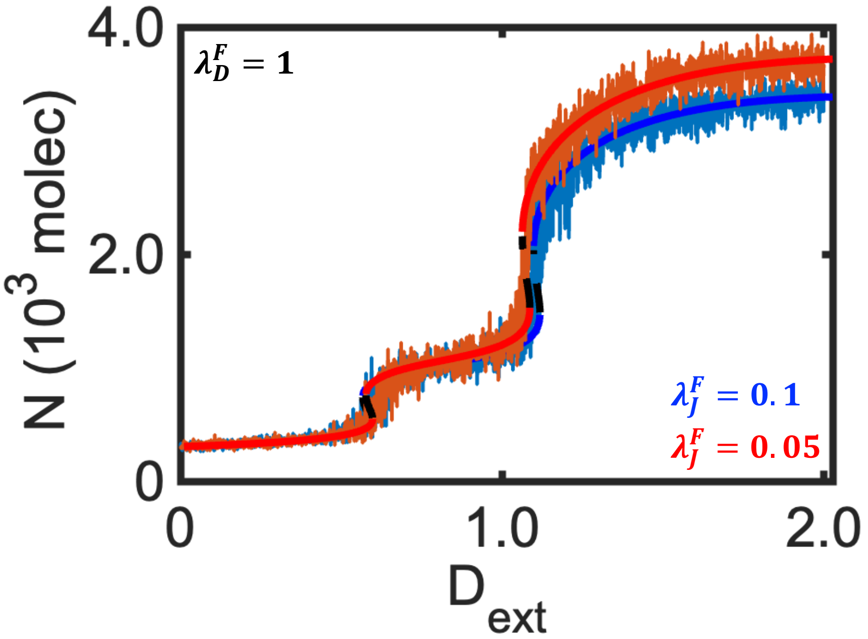

**Figure S10.** Under varied Fringe mediated effect on Notch-Jagged binding ($\lambda_D^F = 1$, $\lambda_J^F < 1$, Case-1), the characteristic behaviour of bifurcation diagram and flickering for Notch (N).

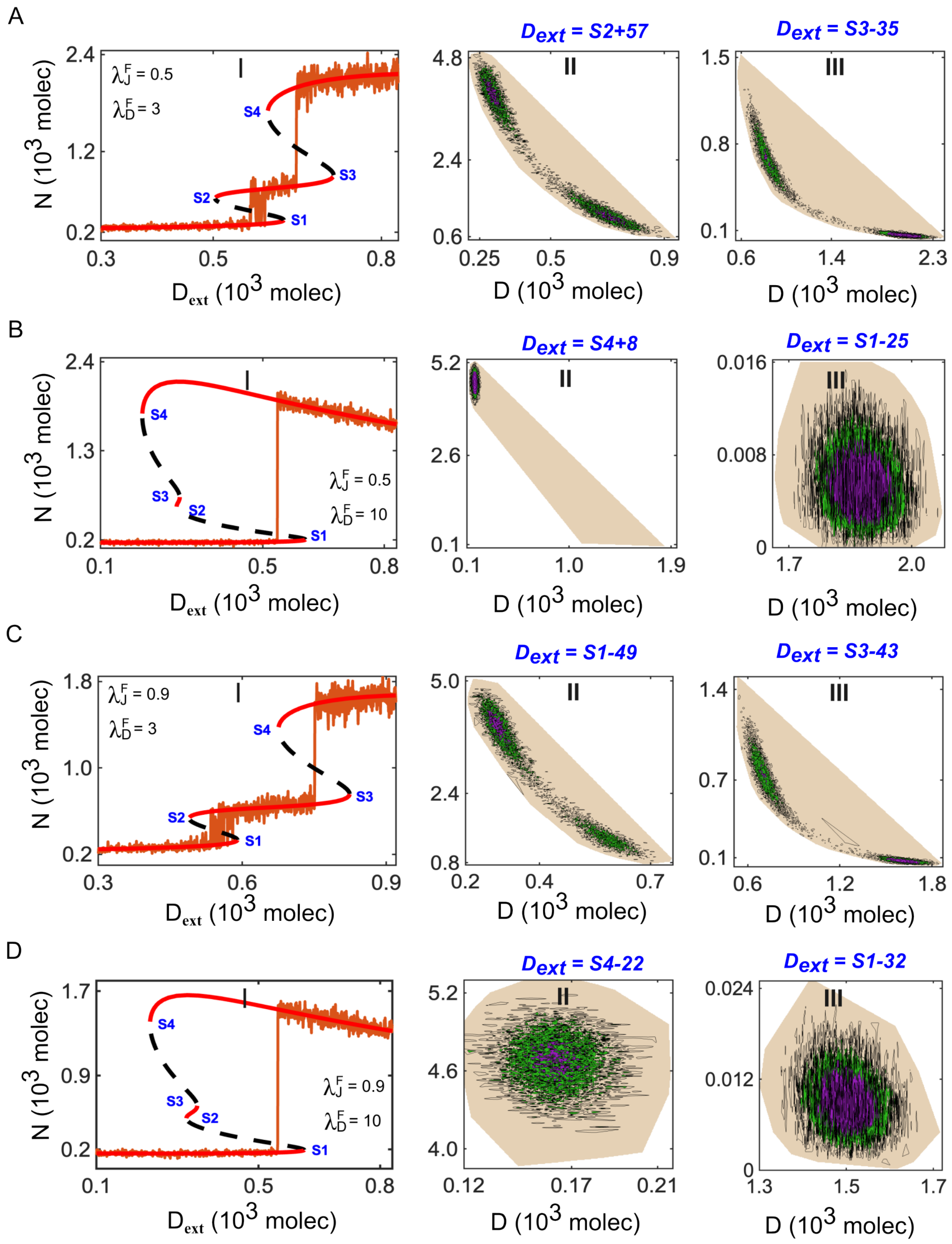

**Figure S11.** Potential landscape of the NDJ circuit system under Fringe mediated effect on both Notch-Delta AND Notch-Jagged binding ($n_j = 5$, WT) ($\lambda_D^F > 1$ and $\lambda_J^F < 1$). (A) Bifurcation diagram exhibiting multiple saddle points ($S1 = 625, S2 = 503, S3 = 715, S4 = 597$) and stochastic potential landscapes under Fringe effect on moderate Delta ($\lambda_D^F = 3$) and Jagged binding ($\lambda_J^F = 0.5$). (B) Bifurcation diagram exhibiting multiple saddle points ($S1 = 605, S2 = 287, S3 = 295, S4 = 202$) and stochastic potential landscapes under Fringe effect on strong Delta ($\lambda_D^F = 10$) and moderate Jagged binding affinity ($\lambda_J^F = 0.5$). (C) Bifurcation diagram exhibiting multiple saddle points ($S1 = 589, S2 = 489, S3 = 823, S4 = 674$) and stochastic potential landscapes under Fringe effect on moderate Delta ($\lambda_D^F = 3$) and strong Jagged binding ($\lambda_J^F = 0.9$). (D) Bifurcation diagram exhibiting multiple saddle points ($S1 = 612, S2 = 321, S3 = 348, S4 = 232$) and stochastic potential landscapes under strong Fringe effect on both Delta ($\lambda_D^F = 10$) and Jagged binding ($\lambda_J^F = 0.9$).

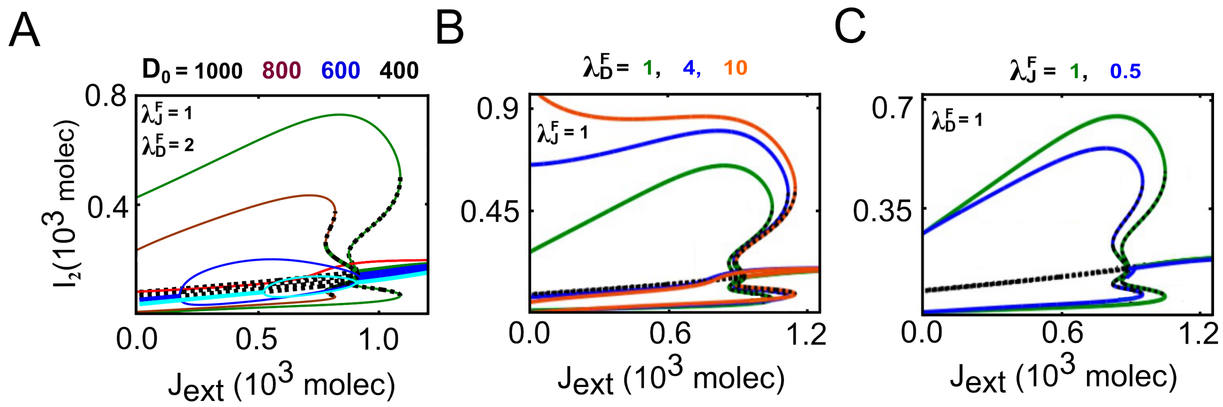

**Figure S12.** Changes in the topology of the two-parameter bifurcation diagram under the moderate effect of Fringe on Delta binding, with (A) and without (B) varying the production rate of Delta ($D_0$), and (C) under the moderate effect of Fringe on jagged binding. The solid (red) lines indicate stable steady states, and the dotted (black) line indicates the unstable steady state.

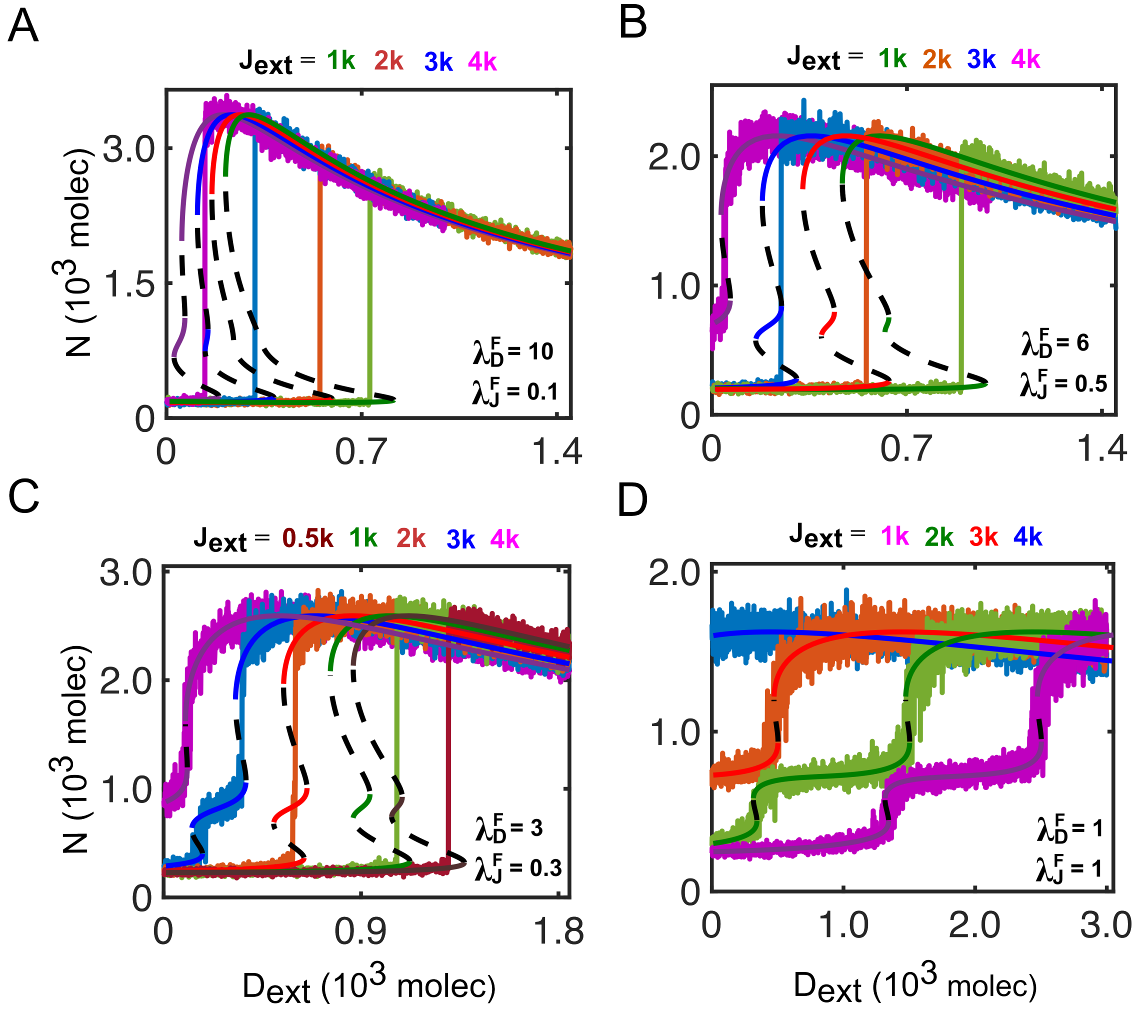

**Figure S13.** Under varied Fringe mediated effect on Notch-Jagged binding ($\lambda_D^F > 1$, $\lambda_J^F < 1$), the characteristic behaviour of bifurcation diagram and flickering for Notch (N) with varying $J_{ext}$.

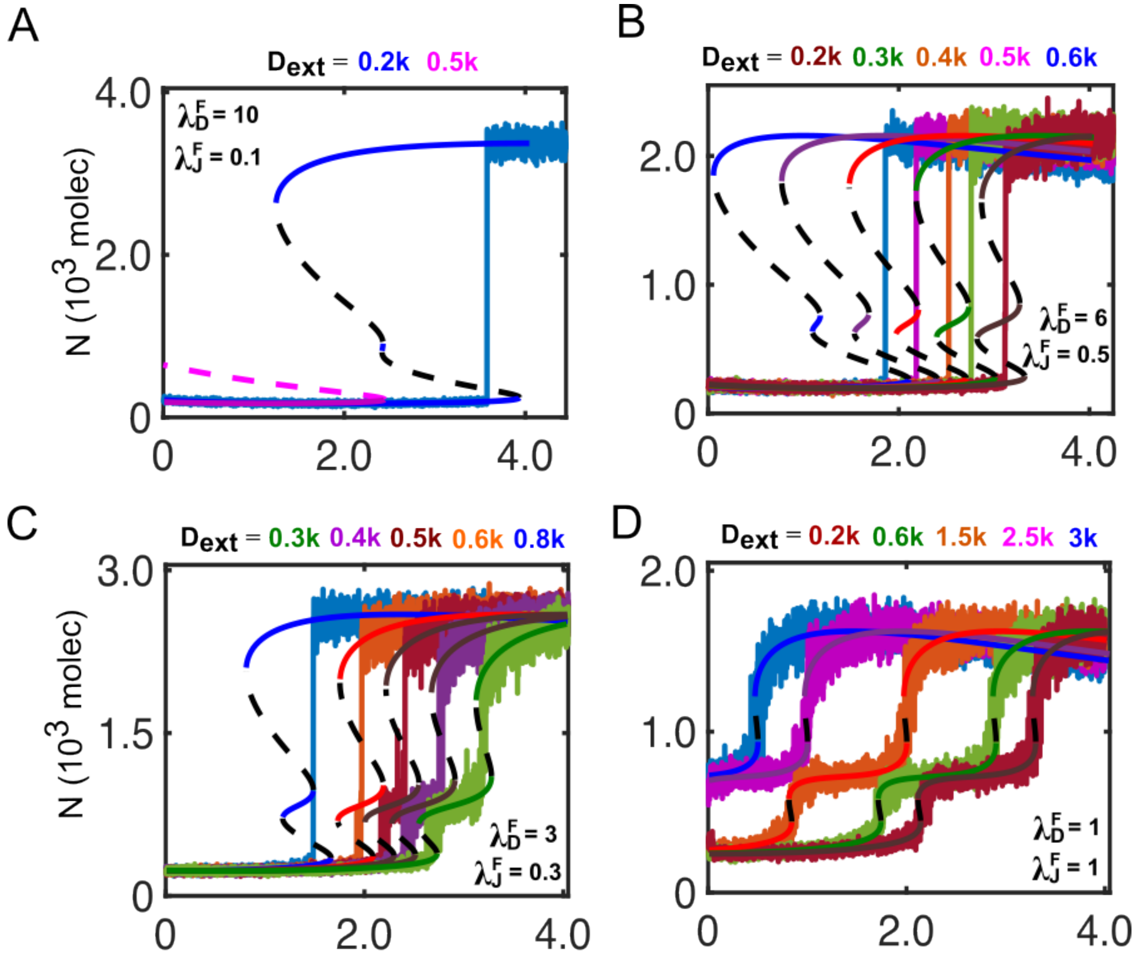

**Figure S14.** Under varied Fringe mediated effect on Notch-Jagged binding ($\lambda_D^F > 1$, $\lambda_J^F < 1$), the characteristic behaviour of bifurcation diagram and flickering for Notch (N) with varying $D_{ext}$.
